# A consensus molecular classification of muscle-invasive bladder cancer

**DOI:** 10.1101/488460

**Authors:** Aurélie Kamoun, Aurélien de Reyniès, Yves Allory, Gottfrid Sjödahl, A. Gordon Robertson, Roland Seiler, Katherine A. Hoadley, Hikmat Al-Ahmadie, Woonyoung Choi, Clarice S. Groeneveld, Mauro A. A. Castro, Jacqueline Fontugne, Pontus Eriksson, Qianxing Mo, Jordan Kardos, Alexandre Zlotta, Arndt Hartmann, Colin P. Dinney, Joaquim Bellmunt, Thomas Powles, Núria Malats, Keith S. Chan, William Y. Kim, David J. McConkey, Peter C. Black, Lars Dyrskjøt, Mattias Höglund, Seth P. Lerner, Francisco X. Real, François Radvanyi, The Bladder Cancer Molecular Taxonomy Group

**Affiliations:** Cartes d’Identité des Tumeurs Program, French League Against Cancer, Paris, France; Department of Pathology, Institut Curie Hospital Group, Paris, France; Division of Urological Research, Department of Translational Medicine, Lund University, Skåne University Hospital Malmö, Sweden; Canada’s Michael Smith Genome Sciences Center, BC Cancer Agency, Vancouver, Canada; Department of Urology, Bern University Hospital, Switzerland; Department of Genetics, Lineberger Comprehensive Cancer Center, University of North Carolina at Chapel Hill, Chapel Hill, NC, USA; Department of Pathology, Memorial Sloan Kettering Cancer Center, New York, NY, USA; Johns Hopkins Greenberg Bladder Cancer Institute and Brady Urological Institute, Johns Hopkins University, Baltimore, MD, USA; Bioinformatics and Systems Biology Laboratory, Federal University of Paraná, Polytechnic Center, Curitiba, Brazil; Division of Oncology and Pathology, Department of Clinical Sciences, Lund University, Lund, Sweden; Department of Medicine, Baylor College of Medicine, Houston, TX, USA; Department of Surgery, Division of Urology, University of Toronto, Mount Sinai Hospital and University Health Network, Toronto, ON, Canada; Institute of Pathology, University Erlangen-Nürnberg, Krankenhausstr 8-10, Erlangen, Germany; Department of Urology and Department of Cancer Biology, University of Texas MD Anderson Cancer Center, Houston, TX, USA; Bladder Cancer Center, Dana-Farber/Brigham and Women’s Cancer Center, Harvard Medical School, Boston, MA, USA; Barts Cancer Institute ECMC, Barts Health and the Royal Free NHS Trust, Queen Mary University of London, London, UK; Genetic and Molecular Epidemiology Group, Spanish National Cancer Research Centre (CNIO), CIBERONC, Madrid, Spain; Molecular & Cellular Biology/Scott Department of Urology, Baylor College of Medicine, One Baylor Plaza, Houston, TX, USA; Department of Genetics, Department of Medicine, Lineberger Comprehensive Cancer Center, University of North Carolina at Chapel Hill, Chapel Hill, NC, USA; Department of Urologic Sciences, University of British Columbia, Vancouver, British Columbia, Canada; Department of Molecular Medicine, Aarhus University Hospital, Aarhus 8200, Denmark; Scott Department of Urology, Dan L. Duncan Cancer Center, Baylor College of Medicine, Houston, TX, USA; Epithelial Carcinogenesis Group, Spanish National Cancer Research Centre (CNIO), CIBERONC, Madrid, Spain; Molecular Oncology, CNRS UMR 144, Institut Curie, Paris, France; Division of Molecular Hematology, Department of Laboratory Medicine, Faculty of Medicine, Lund University, Lund, Sweden; Department of Pathology, Memorial Sloan Kettering Cancer Center, New York, NY 10065, USA; Bladder Cancer Center, Dana-Farber/Brigham and Women’s Cancer Center, Harvard Medical School, Boston, MA, 02215, USA; Oncologie Moléculaire, CNRS UMR 144, Institut Curie, Paris, France; Molecular & Cellular Biology/Scott Department of Urology, Baylor College of Medicine, One Baylor Plaza, Houston, TX 77030, USA; Department of Pathology, The University of Texas MD Anderson Cancer Center, Houston, TX 77030, USA; GenomeDx Biosciences Inc., Vancouver, BC Canada; Cartes d’Identité des Tumeurs Program, Ligue Nationale Contre le Cancer, 75013 Paris, France; Broad Institute of MIT and Harvard, Cambridge, MA, USA; Department of Medicine, Brigham and Women’s Hospital, Harvard Medical School, Boston, MA 02115, USA; Department of Urology, University of Versailles-Saint-Quentin-en-Yvelines, Foch Hospital, Suresnes, France; Department of Translational Medicine, Lund University, Skåne University Hospital, Malmö, Sweden; Genetic and Molecular Epidemiology Group, Spanish National Cancer Research Centre (CNIO), Madrid, Spain; Epithelial Carcinogenesis Group, Spanish National Cancer Research Centre (CNIO), Madrid, Spain; Department of Genitourinary Medical Oncology, The University of Texas MD Anderson Cancer Center, Houston, TX, USA; Department of Bioinformatics and Computational Biology, The University of Texas MD Anderson Cancer Center, Houston, TX 77030, USA

**Author notes:** these authors contributed equally to this work (Aurélien de Reyniès, Yves Allory, Gottfrid Sjödahl, A. Gordon Robertson).

## Abstract

Muscle-Invasive Bladder Cancer (MIBC) is a molecularly diverse disease with heterogeneous clinical outcomes. Several molecular classifications have been proposed, yielding diverse sets of subtypes. This diversity hampers the clinical application of such knowledge. Here, we report the results of a large international effort to reach a consensus on MIBC molecular subtypes. Using 1750 MIBC transcriptomes and a network-based analysis of six independent MIBC classification systems, we identified a consensus set of six molecular classes: Luminal Papillary (24%), Luminal Non-Specified (8%), Luminal Unstable (15%), Stroma-rich (15%), Basal/Squamous (35%), and Neuroendocrine-like (3%). These consensus classes differ regarding underlying oncogenic mechanisms, infiltration by immune and stromal cells, and histological and clinical characteristics. This consensus system offers a robust framework that will enable testing and validating predictive biomarkers in future clinical trials.

## Introduction

Bladder cancer is one of the most frequently diagnosed cancers in North America and Europe (4^th^ in men and 9^th^ in women). Most bladder cancers are urothelial carcinomas, and are classified as either non-muscle-invasive bladder cancer (NMIBC) or muscle-invasive bladder cancer (MIBC), owing to implications for patient management. MIBC is usually diagnosed *de novo*, but may arise from the 10 to 20% of NMIBC cases that eventually progress. MIBC is the most aggressive disease state, and is associated with a five-year survival rate of 60% for patients with localized disease, and less than 10% for patients with distant metastases.

At the molecular level, MIBC is a heterogeneous disease that is characterized by genomic instability and a high mutation rate. Many chromosomal rearrangements and more than 50 oncogenes and tumor suppressor genes have been identified as recurrently altered (Knowles and Hurst, 2015; Robertson et al., 2017). Transcriptome profiling facilitates classifying bladder cancer into molecular subtypes in order to more precisely stratify a patient’s cancer according to prognosis and therapeutic options. Various teams have reported molecular classification of bladder cancers, and several expression-based schemes have been proposed, either considering the full spectrum of non-metastatic bladder cancers (Blaveri et al., 2005; Dyrskjøt et al., 2003; Lindgren et al., 2010; Sjödahl et al., 2012; Tan et al., 2019; Volkmer et al., 2012) or focusing separately on MIBC (Cancer Genome Atlas Research Network, 2014; Choi et al., 2014; Damrauer et al., 2014; Marzouka et al., 2018; Mo et al., 2018; Rebouissou et al., 2014; Robertson et al., 2017; Sjödahl et al., 2017) or on NMIBC (Hedegaard et al., 2016; Hurst et al., 2017). These classifications have considerably advanced our understanding of bladder cancer biology; for example, the association between molecular subtypes and urothelial differentiation, and similarities between subtypes in bladder cancer and other cancers. In addition, specific genomic alterations were found to be enriched in particular molecular subtypes, including mutations targeting genes involved in cell cycle regulation, chromatin remodelling, and receptor tyrosine kinase signaling. Importantly, several reports have highlighted the clinical importance of MIBC molecular stratification, suggesting that responses to chemotherapy and immunotherapy may be enriched in specific MIBC subtypes (Choi et al., 2014; Mariathasan et al., 2018; Rosenberg et al., 2016; Seiler et al., 2017).

The published MIBC classifications share many characteristics, including subtype-specific molecular features; however, the classifications are diverse, containing 2 to 7 molecular subtypes, and using different subtype names. This diversity has hampered transferring subtyping into clinical practice, and highlights that identifying a single set of consensus molecular subtypes would facilitate work to achieve such a transfer.

Here, we report the results of an international collaborative effort to reconcile molecular MIBC classifications. By analyzing six previously published classification schemes and combining public transcriptome data for 1750 tumors, we established a six-class, consensus molecular classification for MIBC. We characterized the consensus classes using additional molecular, pathological and clinical data. To support the use of this consensus molecular classification, we provide a transcriptomic classifier that assigns consensus class labels to single tumor samples (https://github.com/cit-bioinfo/consensusMIBC).

## Results

### Published molecular classifications of MIBC converge on six classes

We used six published MIBC molecular classifications to define a unified consensus subtyping system. We refer to these input classifications as Baylor (Mo et al., 2018), UNC (Damrauer et al., 2014), CIT-Curie (Rebouissou et al., 2014), MDA (Choi et al., 2014), Lund (Marzouka et al., 2018), and TCGA (Robertson et al., 2017). Following the approach outlined in Figure 1, we selected 18 MIBC mRNA datasets (n=1750, Table S1), and assigned each sample to a subtype from each of the six classification systems (Figure S1).

**Figure 1:**
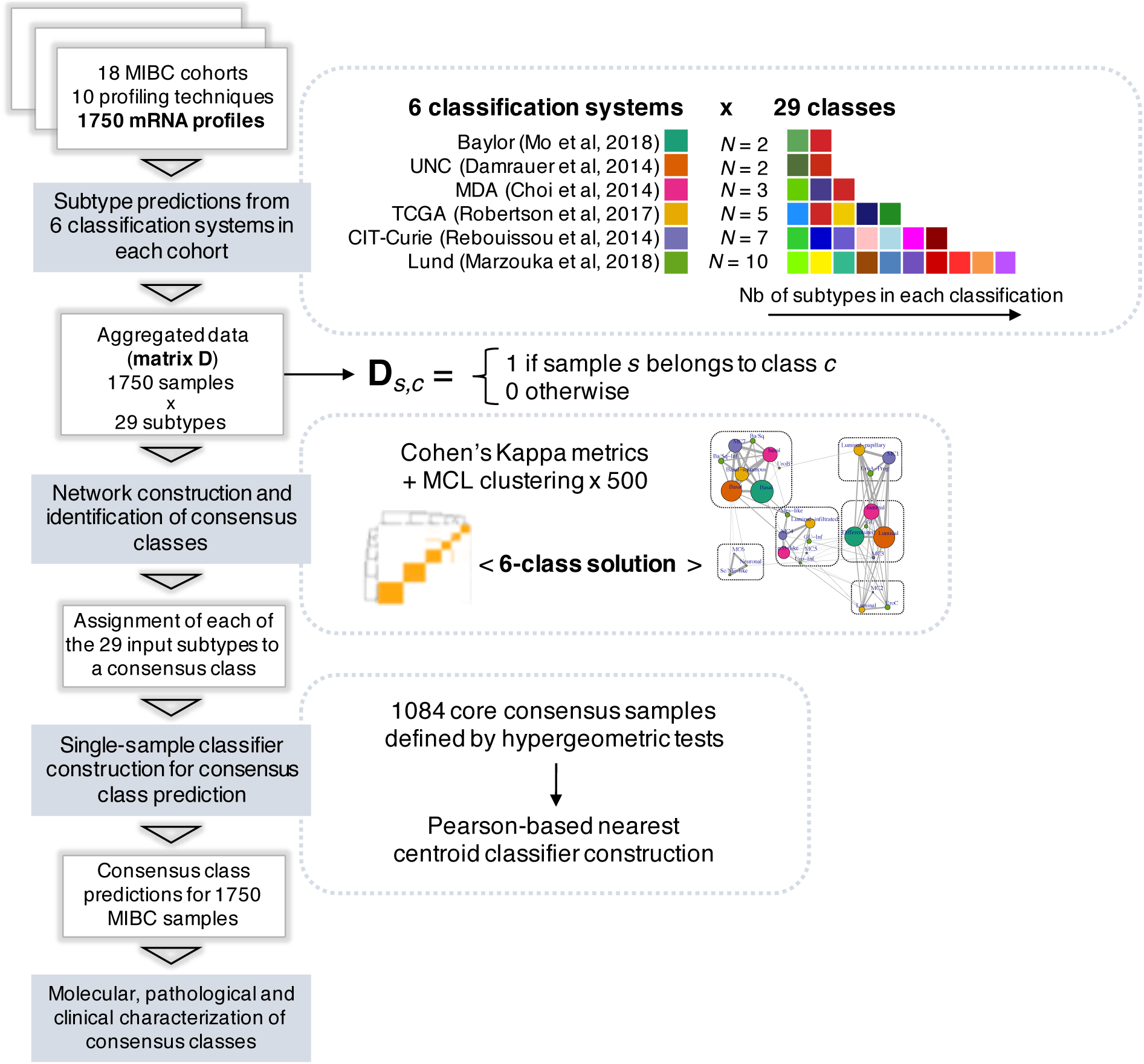
Analytical workflow. We used mRNA classifiers provided by the 6 teams involved in previously published classification systems to subtype 1750 mRNA profiles from 18 independent MIBC cohorts (see Table S1 and Figure S1). A total number of 29 subtypes were considered when summing all classification systems. Using the subtyping results, we could build a 1750 × 29 binary matrix D where a sample “s” was given a value of 1 if assigned to the subtype “m”, and 0 otherwise. The matrix D was used to build a network interconnecting the 29 distinct subtypes. Edges between two subtypes were weighted using a Cohen’s Kappa metric. We performed MCL clustering (Van Dongen, 2008) on this network with 500 bootstrap iterations for several values of inflation factors and used stability scores as weights to calculate weighted silhouette width for each resulting cluster. We used the mean weighted silhouette width as a performance measure to select an inflation factor yielding a robust consensus clustering solution (see Figure S2). An optimal consensus solution was reached for 6 consensus classes. We identified set of 1084 ‘core’ consensus samples, i.e. samples whose initial subtype labels were significantly representative of the consensus classes (*P* < 0.001, hypergeometric test). We used these 1084 core samples to build a nearest-centroid, single-sample classifier based on Pearson’s correlation coefficient, then used the resulting classifier to predict consensus classes on all 1750 MIBC samples. We further characterized the consensus classes using molecular, histological and clinical data.

We built a weighted network of these input subtypes, using Cohen’s Kappa metric to quantify similarities between subtypes from different classification systems, and applied a Markov cluster clustering algorithm (MCL) to identify robust network substructures corresponding to potential consensus classes (Methods, Figure S2). We identified a 6-cluster solution, defining six biologically relevant consensus molecular classes, which we labelled as: Luminal Papillary (LumP), Luminal Non-Specified (LumNS), Luminal Unstable (LumU), Stroma-rich, Basal/Squamous (Ba/Sq), and Neuroendocrine-like (NE-like) (Figure 2A). Considerations motivating our choices for these consensus names are detailed in the Supplementary Note.

**Figure 2:**
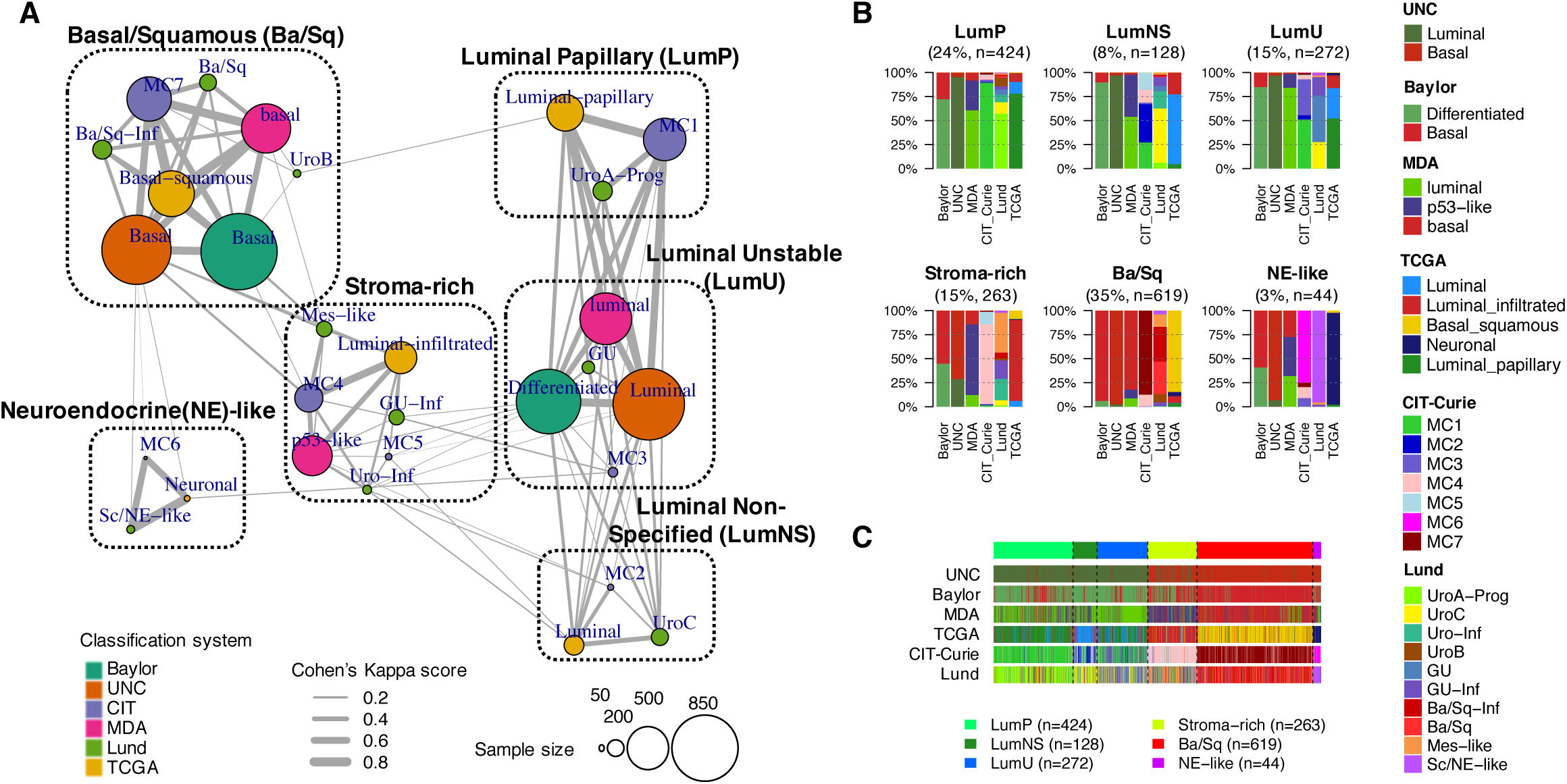
The six consensus classes and their relation to input molecular subtypes. (**A**) Clustered network by MCL clustering. The 6-consensus class solution obtained with MCL clustering on the Cohen’s Kappa-weighted network is represented by the 6 cliques surrounded by black dotted rectangles (see Supplementary Note for the naming of consensus classes) The circles inside each clique symbolize the input subtypes associated with each consensus class and are colored according to their matching classification system. Circle size is proportional to the number of samples assigned to the subtype. Edge width between subtypes is proportional to the Cohen’s Kappa score, which assess the level of agreement between two classification schemes. (**B**) Input subtypes repartitioned among each consensus class. Consensus classes were predicted on 1750 MIBC samples using the single-sample classifier described in Methods (see Table S2, Figure S3). Here, the samples are grouped by their predicted consensus class label: LumP, LumN, LumU, Stroma-rich, Ba/Sq and Neuroendocrine (NE)-like. For each consensus class, a barplot shows the proportion of samples assigned in each input subtype of each input classification system. See also Figure S3 for additional visualizations of consensus classes distributions across input subtypes and across datasets. (**C**) Relationship between subtyping results from the six input classification schemes. Samples are ordered by predicted consensus classes.

The network of consensus classes also revealed a core set of consensus samples (Methods), i.e., tumor samples representative of each consensus class on the basis of their initial subtyping by the 6 classification systems. We used these core samples to build a single-sample transcriptomic classifier. The classifier was trained on approximately one third of these samples (n=403), and achieved a 97% mean balanced accuracy on the remaining two thirds of core samples (n=681) (Figure S3).

The six molecular classes had variable sample sizes, with Ba/Sq and LumP being the most prevalent (35% and 24% of all samples, respectively). The remaining 41% of samples were LumU (15%), Stroma-rich (15%), LumNS (8%), and NE-like (3%) tumors (Figure 2B). The consensus classification was strongly associated with each of the initial classification systems (Chi-square *P*<10^−165^) (Figures 2, S4).

We also compared the consensus classes to the TCGA pan-cancer integrative classification (Hoadley et al., 2018) (Figure S4). We observed enrichments between the Ba/Sq consensus class and the squamous cell carcinoma C27:Pan-SCC pan-cancer cluster (*P*=1.1×10^−11^), and between the Stroma-rich class and the stroma-driven C20:Mixed (Stromal/Immune) pan-cancer cluster (*P*<2.2×10^−16^).

### Transcriptomic characterization of the six consensus molecular classes

We used mRNA data from all 1750 samples to characterize consensus classes with molecular gene signatures for bladder cancer pathways and for tumor microenvironment infiltration (Figure 3, Table S3).

**Figure 3:**
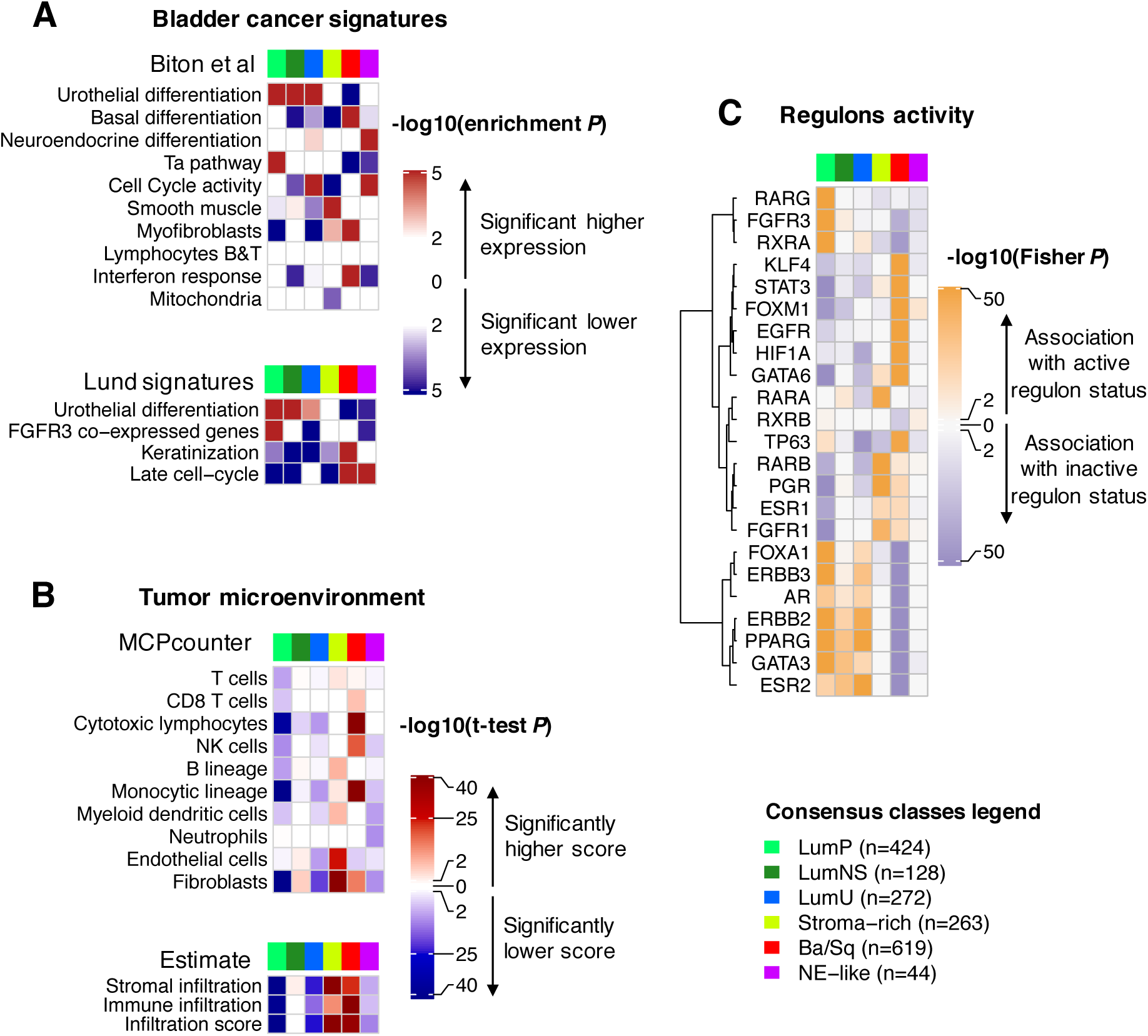
Characterization of tumor and stroma signals using published mRNA signatures and regulon analysis. Description of gene sets and detailed statistics is available in Table S3 (**A**) We performed a Gene Set Analysis (GSA (Efron and Tibshirani, 2007)) in each dataset to test the significance of differential expression of specific bladder cancer related signatures in each consensus class compared to the others. The heatmaps show Stouffer combined GSA p-values over all datasets. The upper panel refers to bladder cancer gene sets extracted from the ICA components described in Biton *et* al (Biton et al., 2014) (see Methods). The lower panel displays other bladder cancer-specific signatures retrieved from the literature: urothelial differentiation, keratinization and late cell-cycle signatures from Eriksson *et al* (Eriksson et al., 2015), and an *FGFR3* co-expressed signature from Sjödahl *et al* (Sjödahl et al., 2012). (**B**) We used two mRNA-based computational tools to characterize tumor microenvironment. The characterization includes an estimate of microenvironment immune and stromal cell subpopulations using MCPcounter (Becht et al., 2016) and a more global measure of stromal and immune infiltrates by ESTIMATE (Yoshihara et al., 2013). We ran MCPcounter and ESTIMATE independently on each dataset and used t-tests to compare scores for each consensus class relative to the others. The heatmaps show Stouffer combined t-test p-values over all datasets. **(C)** We computed discrete regulon status (1 for active regulon status, 0 for neutral status, −1 for inactive regulon status) in each dataset, as described in Methods and in (Robertson et al., 2017). We evaluated the association between each regulon status and each consensus class using Fisher exact tests; the heatmap illustrates the resulting p-values.

Differentiation-associated mRNA signatures were strongly associated with the consensus classes. Tumors from the three luminal classes overexpressed urothelial differentiation signatures (*P*<2.0×10^−4^), including the PPARG/GATA3/FOXA1-related Lund signature (Eriksson et al., 2015). In contrast, Ba/Sq and NE-like tumors overexpressed gene signatures associated with basal (*P*=2.1×10^−25^) and neuroendocrine differentiation (*P*=3.2×10^−26^), respectively.

In addition to their urothelial differentiation status, the three luminal classes exhibited distinct molecular signatures. LumP tumors were characterized by high expression of a non-invasive Ta pathway signature (Biton et al., 2014) (*P*=6.9×10^−199^) and were strongly associated with FGFR3 transcriptional activity as measured by an FGFR3 co-expressed gene signature (Sjödahl et al., 2012) (*P*=2.5×10^−95^). LumNS tumors displayed elevated stromal infiltration signatures, mainly fibroblastic, compared to the other luminal tumors (*P*=6.4×10^−8^). LumU tumors had a higher cell cycle activity than the other luminal tumors (*P*=6.4×10^−8^).

Stroma-rich samples displayed intermediate levels of urothelial differentiation. They were mainly characterized by stromal infiltration as summarized by ESTIMATE (Yoshihara et al., 2013) stromal scores, with a specific overexpression of smooth muscle (*P*=2.8×10^−143^), endothelial (*P*=2.1×10^−25^), fibroblast (*P*=1.4×10^−76^), and myofibroblast (*P*=8.0×10^−4^) gene signatures.

Immune infiltration was mainly found within Ba/Sq and Stroma-rich tumors, but the two classes were characterized by distinct immune cell populations, as measured by MCPcounter signatures (Becht et al., 2016). Ba/Sq tumors were enriched in cytotoxic lymphocytes (*P*=9.8×10^−48^) and NK cells (*P*=2.2×10^−20^), whereas Stroma-rich tumors overexpressed T cell (*P*=1.6×10^−5^) and B cell markers (*P*=4.9×10^−11^). LumNS tumors were the only luminal type associated with any immune infiltration signals, mainly B (*P*=1.7×10^−3^) and T (*P*=4.4×10^−3^) lymphocytes. We detected no transcriptomic markers of immune infiltration in NE-like tumors. Estimation of tumor purity with ABSOLUTE (Carter et al., 2012) in the TCGA samples confirmed that Stroma-rich and Ba/Sq tumors contained higher levels of non-tumor cells (Figure S5).

Analyses of regulatory units (i.e. regulons) for 23 regulator genes previously reported as associated with bladder cancer (Castro et al., 2016; Robertson et al., 2017) were consistent with the mRNA signatures assessed (Figure 3C). Luminal tumors, which overexpressed strong urothelial differentiation signals, were associated with active *PPARG* and *GATA3* regulons (*P*<5.3×10^−32^; *P*<7.6×10^−21^). *FGFR3* regulon activity was specifically associated with LumP tumors (*P*=1.2×10^−94^), and Ba/Sq tumors showed a strong association with *STAT3* regulon activity (*P=*4.2×10^−109^), consistent with their expressing a keratinization gene signature (Eriksson et al., 2015). Additionally, the regulon analysis showed an association of *HIF1A* regulon activity with Ba/Sq tumors (*P*=3.7×10^−94^), suggesting that this class is associated with a hypoxic microenvironment. *EGFR* regulon activity was also specifically associated with Ba/Sq tumors (*P*=1.4×10^−77^), consistent with previously reported findings (Rebouissou et al., 2014).

### Genomic alterations associated with the consensus molecular classes

We used TCGA exome data to identify class-specific mutations (Figure 4A, Table S4), and ran GISTIC2 (Mermel et al., 2011) on 600 available copy number profiles, grouped by consensus class, to identify class-specific copy number aberrations (CNA) (Table S5). In addition, we combined all CNA, gene fusion, and gene mutation data from the 18 cohorts to generate comprehensive profiles of genomic alterations of seven key bladder cancer genes (*FGFR3, CDKN2A, PPARG, ERBB2, E2F3, TP53* and *RB1*) according to consensus class (Figure 4B).

**Figure 4:**
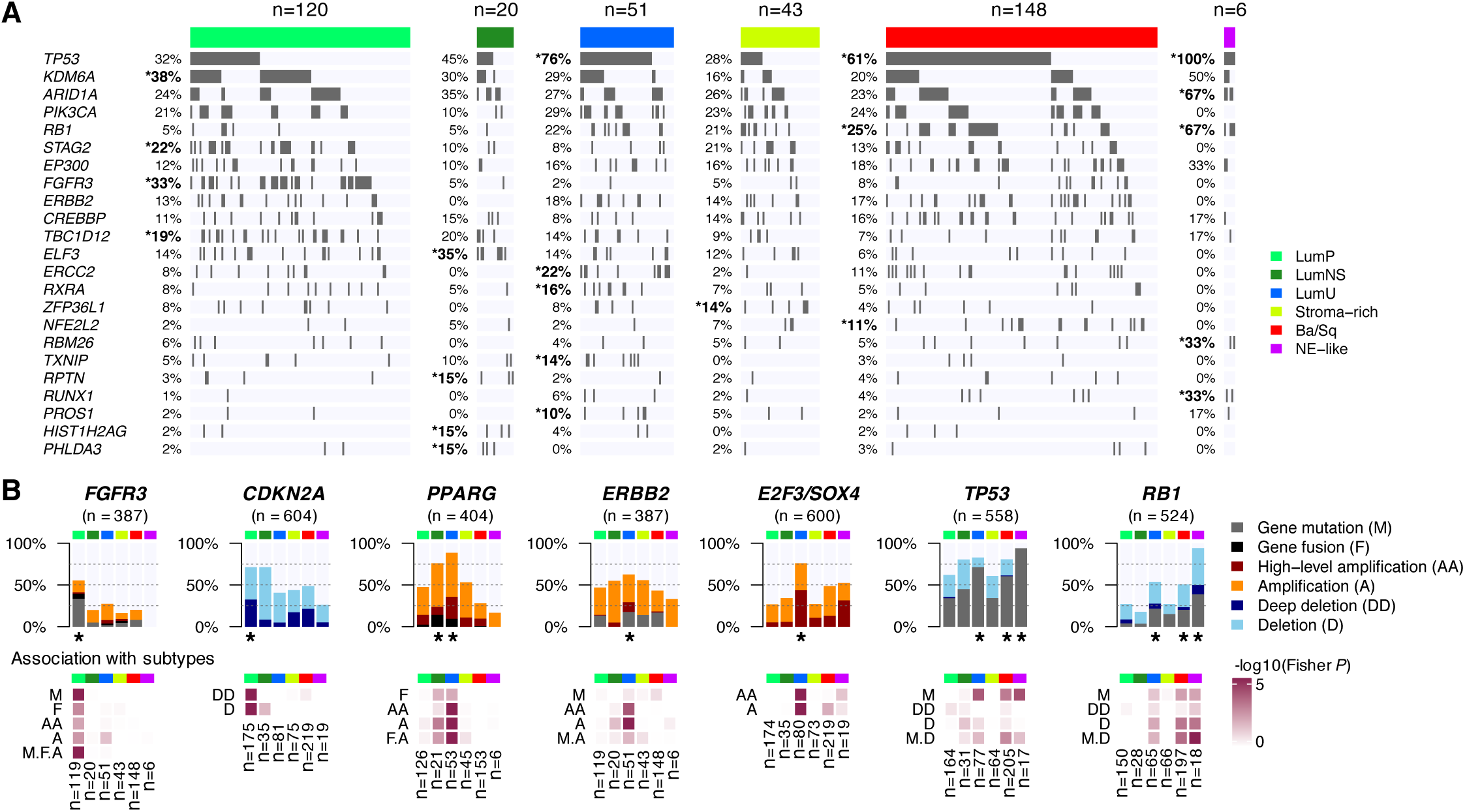
Genomic alterations associated with consensus classes. (**A**) We used the available exome data from 388 TCGA samples to study the association between consensus classes and specific gene mutations (see Table S4 and Figure S5). The panel displays the 23 genes with MutSig p-values<0.001 that were either found in ≥10% of all tumors, or significantly overrepresented within one of the consensus classes (Fisher *P* <0.05 and frequency within a consensus class >10%). Gene mutations that were significantly enriched in one consensus class are marked by an asterisk. (**B**) Combined genomic alterations associated with seven bladder cancer-associated genes and statistical association with consensus classes. Upper panels: Main alteration types after aggregating CNA profiles (see Table S5) from CIT (n=87), Iyer (n=58), Sjödahl (n=29), Stransky (n=22), and TCGA (n=404) data; exome profiles (n=388) and *FGFR3* and *PPARG* fusion data (n=404) from TCGA data; *CDKN2A* and *RB1* MLPA data from CIT (n=86; n=85) and Stransky (n=16; n=13) data; *FGFR3* mutation data from MDA (n=66), CIT (n=87), Iyer (n=39), Sjödahl (n=28), and Stransky (n=35); *TP53* mutation data from MDA (n=66), CIT (n=87), Iyer (n=39), Sjödahl (n=28), and Stransky (n=19); and *RB1* mutation data from MDA (n=66), CIT (n=85), Iyer (n=39) and Stransky (n=13). Lower panels: Associations between each consensus class, each type of gene alteration, and the combined alterations were evaluated by Fisher’s exact test. Consensus classes significantly enriched with alterations of these candidate genes are marked with a black asterisk.

LumP tumors were enriched in *FGFR3* (*P*=1.4×10^−11^), *KDM6A* (*P*=2.2×10^−03^), and *STAG2* mutations (*P*=0.012). Adding mutation data from *FGFR3* targeted sequencing in additional cohorts (n=255), the proportion of *FGFR3*-mutated LumP tumors reached 40% (*P*=1.6×10^−23^). Assembling mutations, fusions, and copy number amplifications, *FGFR3* alterations were enriched in LumP tumors (*P*=1.9×10^−11^). *CDKN2A* MLPA (Multiplex Ligation-dependent Probe Amplification) and CNA data for 604 tumors revealed 33% of *CDKN2A* homozygous/deep deletions in LumP tumors, corresponding to a strong enrichment as compared to other classes (*P*=3.8×10^−8^). These deletions were consistent with the enrichment of LumP tumors within the TCGA pan-cancer iCluster C7:Mixed(Chr9 del) (*P*=1.6×10^−10^), which is characterized by Chr 9 deletions (Figure S4).

The LumNS class was mainly characterized by an enrichment of mutations in *ELF3* (35%, *P*=0.004), which is an early regulator of urothelial differentiation and is activated by PPARG (Böck et al., 2014). *PPARG* was significantly altered as well, with 76% of LumNS tumors harboring either amplifications or fusions (*P*=5.7×10^−3^).

LumU tumors also harbored frequent *PPARG* alterations (89%, *P*=1.9×10^−11^) and high-level amplifications of a 6p22.3 region that contains *E2F3* and *SOX4* (76%, *P*=3.0×10^−12^). *ERBB2* amplifications were overrepresented in LumU tumors (*P*=4.3×10^−8^), but no significant association was found between *ERBB2* mutations and any of the consensus classes. In contrast with the other luminal classes, LumU tumors were associated with mutations in *TP53* (76%, *P*=3.4×10^−5^) and in *ERCC2*, which codes for a core nucleotide-excision repair component (22%, *P*=0.006). More generally, LumU was the most genomically altered class (Figure S5), displaying the highest number of copy number alterations (*P*=1.8×10^−16^), the highest somatic mutation load (*P*=0.009), and including more APOBEC-induced mutations than other consensus classes (*P*=0.01). These features of genomic instability and the association with *ERBB2* amplifications were consistent with the enrichment of LumU tumors within the TCGA pan-cancer subtypes C2:BRCA(HER2 amp) (characterized by frequent *ERBB2* amplifications, *P*=4.0×10^−5^) and C13:Mixed (Chr8 del) (enriched in highly aneuploid tumors, *P*=3.8×10^−9^), shown in Figure S4 (Hoadley et al., 2018).

For Ba/Sq tumors, as shown previously (Choi et al., 2017), the most frequently mutated genes were *TP53* (*P*=5.8×10^−4^), *NFE2L2* (*P*=0.002) and *RB1* (*P*=0.002). Aggregated mutation data revealed that 58% (134/232, *P*=0.009) and 20% (43/224, *P*=0.007) of Ba/Sq tumors contained mutations in *TP53* and *RB1*, respectively; these mutations co-occurred in 14% (32/224) of Ba/Sq cases. Ba/Sq tumors were also strongly associated with genomic deletions of 3p14.2, which occurred in 49% of cases (*P* = 1.5×10^−13^).

When combining all available data on genomic alterations of *TP53* and *RB1*, we observed a strong enrichment of concomitant *TP53* and *RB1* inactivation in NE-like tumors. *TP53* was ubiquitously mutated in these tumors (94%, *P*=9.7×10^−5^), and co-occurred with *RB1* inactivation by either mutations or deletions (94%, *P*=2.2×10^−6^).

### Histological patterns associated with the consensus molecular classes

To characterize the consensus molecular classes from a histological perspective, we assembled annotations for urothelial histological variants and specific morphological patterns (Figure 5 and Figure S6). As expected, Ba/Sq tumors included 79% of tumors in which histological review identified squamous differentiation (126/159, *P*=3.6×10^−32^). However, the Ba/Sq class extended beyond this histological subtype, with only 42% (126/303) of Ba/Sq tumors associated with squamous differentiation as identified by pathologists. Similarly, NE-like tumors were strongly associated with neuroendocrine variant histology, with 72% of histologically reviewed NE-like tumors showing neuroendocrine differentiation (13/18, *P*=9.7×10^−22^). LumP tumors were enriched with papillary morphology as compared to other consensus classes (*P*=1.2×10^−12^). This pattern was observed in 59% (82/139) of histologically reviewed LumP tumors, although it was frequently found in other luminal classes (42% in LumNS and 31% in LumU). LumNS tumors were enriched in micropapillary variant histology (36%, 9/25, *P*=0.001) and were commonly associated with carcinoma *in situ* (CIS) (80%, 4/5, *P*=0.005).

**Figure 5:**
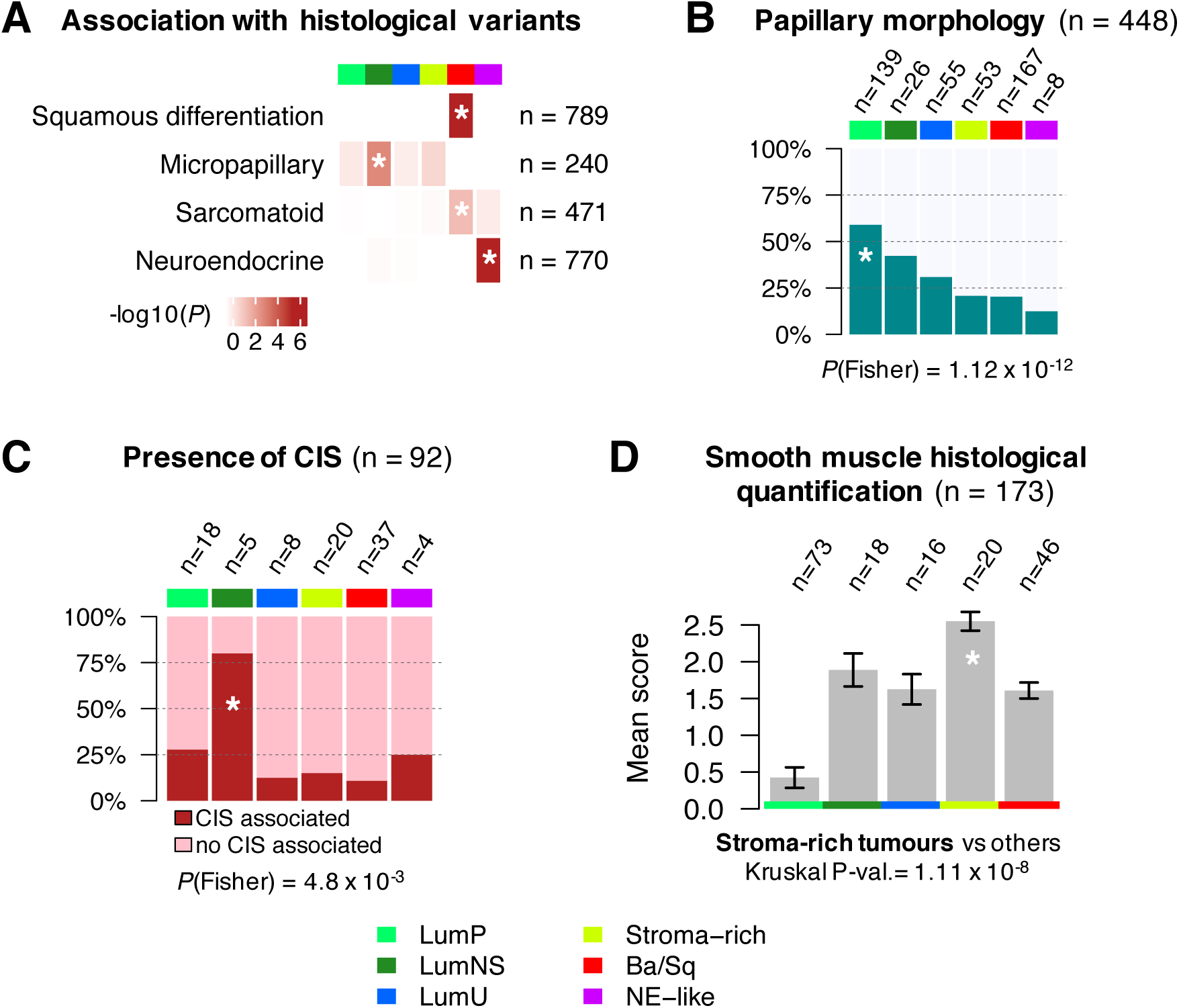
Histopathological associations with consensus classes. (**A**) Histological variant overrepresentation within each consensus class. One-sided Fisher exact tests were performed for each class and histological pattern. Pathological review of histological variants was available for several cohorts: squamous differentiation was evaluated in CIT (n=75), MDA (n=46), Sjödahl2012 (n=23), Sjödahl2017 (n=239), and TCGA (n=406) cohorts; neuroendocrine variants were reviewed in CIT (n=75), MDA (n=46), Sjödahl2017 (n=243), and TCGA (n=406) cohorts; micropapillary variants were reviewed in CIT (n=75), MDA (n=46) and TCGA cohorts (n=118 FFPE tumor slides from TCGA were reviewed by Y.A. and J.F. for this study). Results are displayed on the heatmap as –log_10_(Fisher’s *P*). Detailed sample counts within each class are given on Figure S5. (**B**) Occurrence of papillary morphology in tumors from the TCGA cohort (n=401) and the CIT cohort (n=47). (**C**) Proportion of samples with associated CIS within each consensus class in tumors from the CIT cohort (n=84) and the Dyrskjøt cohort (n=8). (**D**) Smooth muscle infiltration from images for 173 tumor slides from the TCGA cohort. Each sample was assigned a semi-quantitative score ranging from 0 to 3 (0 = absent, 1 = low, 2 = moderate, 3 = high) to quantify the presence of large smooth muscle bundles. The barplot shows means and standard errors for each class.

A pathological review of stromal infiltration in slide images corresponding to the TCGA tumor samples confirmed that Stroma-rich tumors contained a higher proportion of smooth muscle cells (*P*=1.1×10^−8^), consistent with the strong smooth muscle-related mRNA expression characterizing these tumors.

### Association of the consensus molecular classes with clinical characteristics, survival outcomes, and therapeutic opportunities

The consensus classes were associated with gender, stage, and age (Figure 6A). Ba/Sq tumors were overrepresented in females (*P*=1.4×10^−5^) and in higher clinical stages (*P*=2.8×10^−7^). These data confirm previously published results (Damrauer et al., 2014; Rebouissou et al., 2014; Robertson et al., 2017; Sjödahl et al., 2012). The LumP and LumU consensus classes were enriched in T2 vs. T3-4 tumors (*P*=0.009 and *P*=4.2×10^−4^) as compared to other classes. Patients less than 60 years old were overrepresented among LumP tumors (*P*=0.001), whereas the LumNS consensus class was enriched with older patients (> 80 years; *P*=0.03).

**Figure 6:**
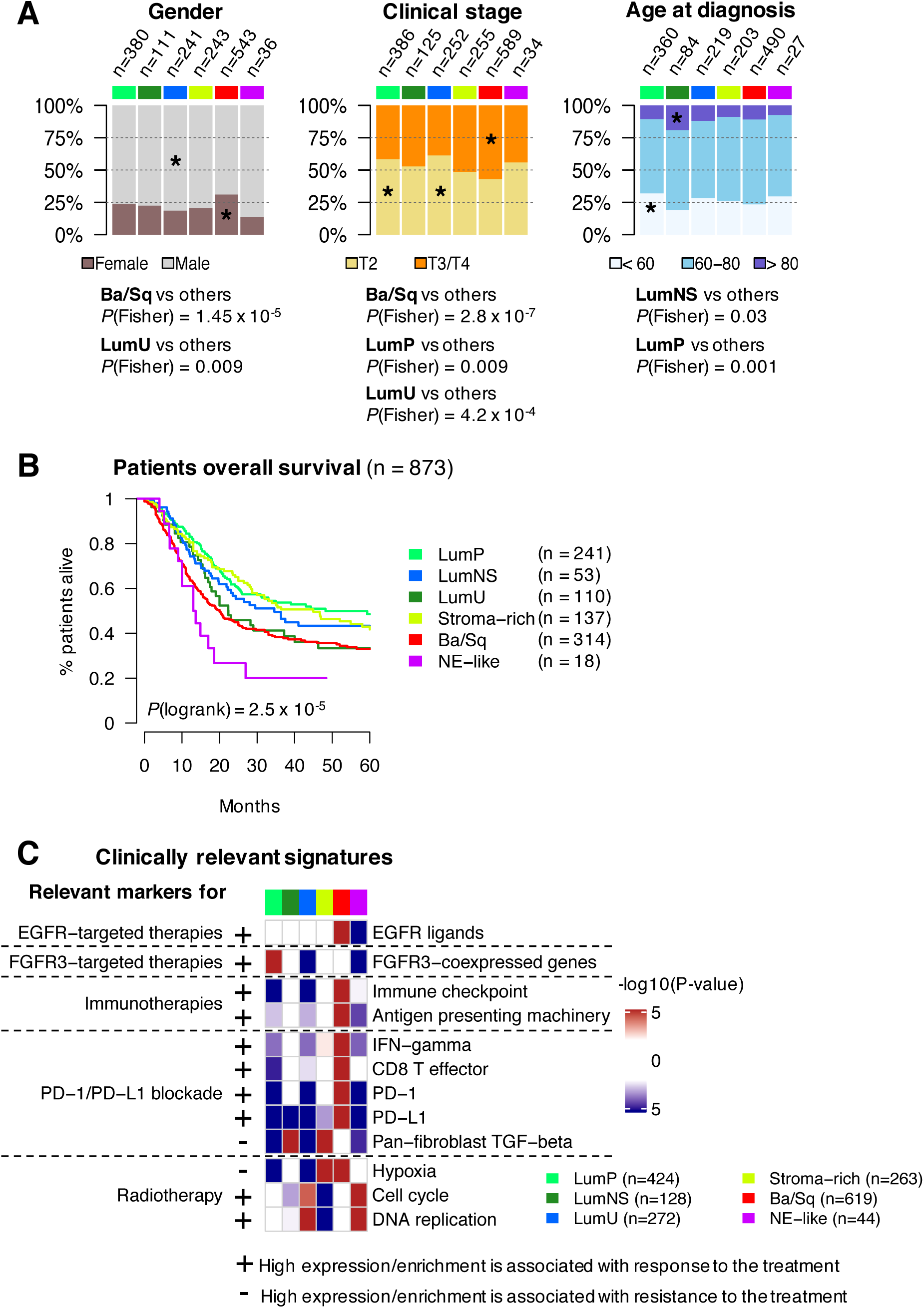
Clinical characteristics and prognostic associations. (**A**) Association of consensus classes with gender (n=1554), clinical stage (n=1641), and age category (n=1378). (**B**) 5-year overall survival stratified by consensus class (see also Figure S7). Kaplan-Meier curves were generated from 873 patients with available follow-up data. Patients who had received neoadjuvant chemotherapy were excluded from the survival analysis. Detailed statistics of univariate and multivariate survival analyses are given in Table S6. (**C**) We selected a set of clinically relevant gene signatures (see Table S7) and performed a Gene Set Analysis (GSA (Efron and Tibshirani, 2007)) in each dataset to test the significance of their differential expression in each consensus class relatively to the others. We used one-sided t-tests to assess the differential expression of single genes (PD-1 and PD-L1). The heatmaps show Stouffer combined p-values over all datasets. Plus/minus annotation of gene sets indicates association of high gene expression levels with response/resistance to the corresponding therapy.

Overall survival was strongly associated with the consensus classes (Figure 6B, *P*=2.5×10^−5^): patients with LumP tumors had the best prognosis. The two other luminal classes were associated with poorer patient prognosis (HR_LumNS/LumP_=1.51, *P*=4.7×10^−^ 2; and HR_LumU/LumP_=1.32, *P*=0.12), although these differences were respectively modest and not significant, in this setting. Patients with stroma-rich tumors showed a similar overall survival to that associated with LumP tumors (HR_Stroma-rich/LumP_=1.18, CI_95_ = [0.85, 1.63]), and their survival was independent the differentiation status of the tumor sample (Figure S7).

Ba/Sq tumors were associated with a poor prognosis (HR_BaSq/LumP_=1.8, *P*=5.7×10^−6^), consistent with previous studies (Rebouissou et al., 2014). Finally, NE-like tumors were associated with the worst prognosis (HR_NE-like/LumP_=2.4, *P*=3.3×10^−3^). Ba/Sq and NE-like consensus classes remained significantly associated with worse overall survival in a multivariate Cox model that combined consensus classes (with the LumP class as reference), TNM, and patient age (respectively *P*=0.002 and *P*=0.05, Table S6B).

We characterized the consensus classes using several clinically relevant mRNA signatures (Figure 6C, Table S7). The FGFR3 signature was strongly and specifically activated in LumP tumors (*P*=6.5×10^−14^), suggesting that FGFR3-targeted therapies warrant investigation in patients with tumors of this consensus class. Ba/Sq tumors expressed high levels of EGFR receptor and its ligands (*P*=2.7×10^−11^), which may be associated with a sensitivity to EGFR-targeted therapies, as suggested by previously reported *in vitro* and *in vivo* experiments (Rebouissou et al., 2014). Ba/Sq tumors also strongly expressed immune checkpoint markers (*P*=2.1×10^−8^) and antigen-presenting machinery genes (*P*=3.0×10^−8^), suggesting that such tumors might be more responsive to immunotherapies. Studies integrating mRNA signatures with data on response to anti-PD1/PD-L1 therapies (Ayers et al., 2017; Mariathasan et al., 2018) have reported associations of anti-PD1/PD-L1 response with high levels of CD8 T cells, high interferon gamma signals, and low activity of the TGF-beta pathway. However, considering this combination of factors, no consensus class had an expression profile that clearly suggested either response or resistance to anti-PD1/PD-L1 therapies. In contrast, NE-like and LumU tumors both had a profile associated with response to radiotherapy (Horsman and Overgaard, 2016; Pawlik and Keyomarsi, 2004), showing elevated cell cycle activity (*P*_*NE-like*_=1.1×10^−10^, *P*_*LumU*_=5.3×10^−5^) and low hypoxia signals (*P*_*NE-like*_=1.0×10^−2^, *P*_*LumU*_=4.6×10^−15^).

Finally, we performed a consensus class-based retrospective analysis of outcome of patients receiving neoadjuvant chemotherapy (Choi et al., 2014; Seiler et al., 2017) (NAC) and patients treated with the anti-PD-L1 antibody Atezolizumab (Mariathasan et al., 2018) (IMvigor210) (Figure 7). Analysis of overall survival and response showed that consensus classes were associated with variable responses to therapy. The results did not suggest any significant association of a consensus class with patient response to NAC. However, we observed an enrichment in Atezolizumab responders among patients with LumNS (*P*=5.0×10^−2^), LumU (*P*=4.4×10^−3^), and NE-like (*P*=1.2×10^−2^) tumors. Particularly, NE-like tumors may identify a subset of good responders to immune checkpoint inhibitors, as suggested by recent results (Kim et al., 2019).

**Figure 7:**
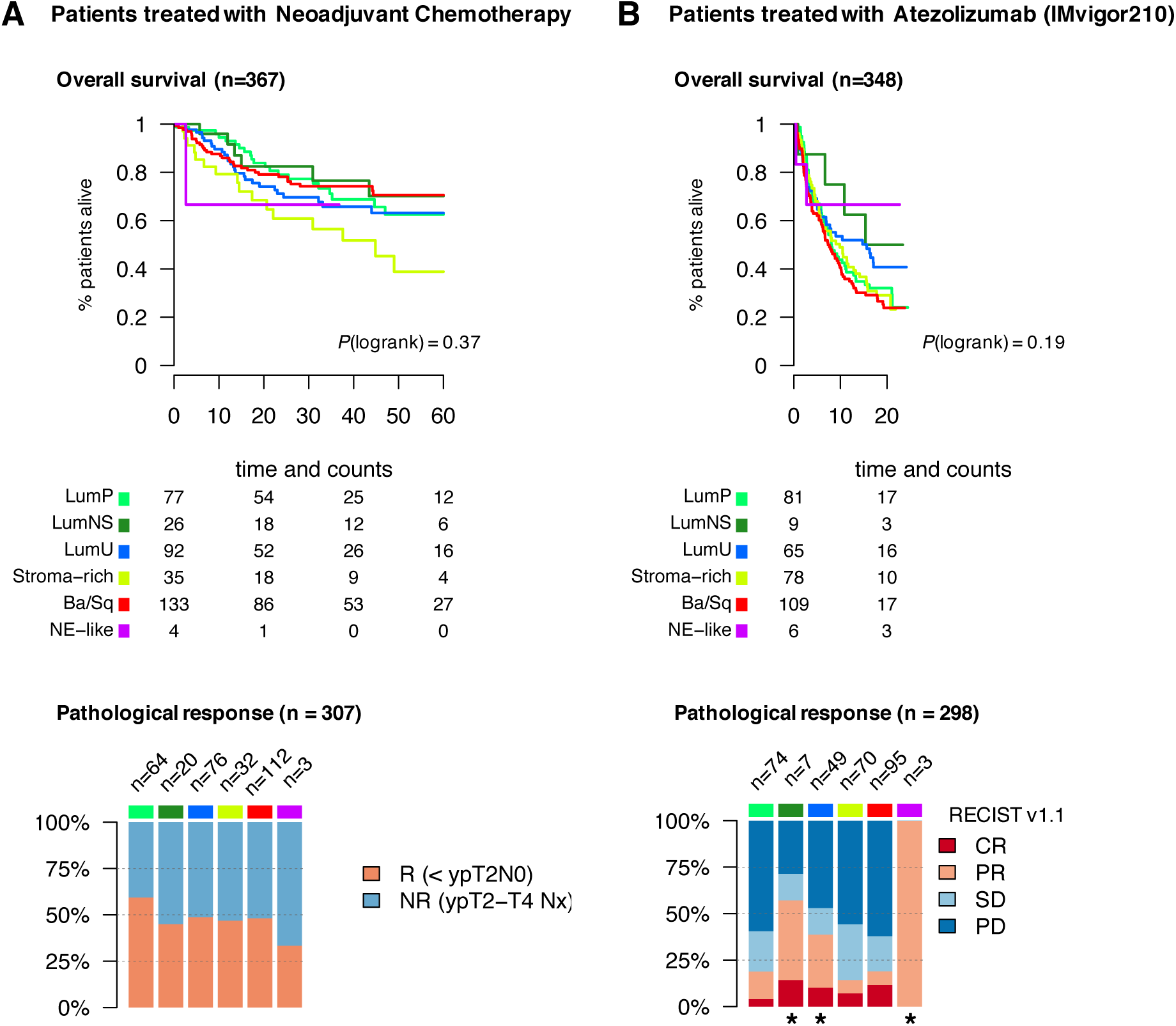
Response to neoadjuvant chemotherapy and PD-L1 blockade. Analysis of overall survival and response data from patients who had received neoadjuvant chemotherapy (NAC) and patients treated with the anti PD-L1 Atezolizumab (Mariathasan et al., 2018) (IMvigor210). The pre-treatment tumor samples from these patients were classified according to the consensus molecular classification. (**A**) Overall survival and response data to neoadjuvant chemotherapy (NAC). Patients treated with NAC were selected from Seiler (n=262), MDA MVAC (n=22, GSE70691), MDA DDMVAC (n=33, GSE69795), and Aarhus.3 (n=50, Christensen et al, to be published in Journal of Clinical Oncology) cohorts. Pathological response to NAC was obtained from MDA MVAC (n=23), MDA DDMVAC (n=34, GSE69795), Seiler (n=199), and Aarhus.3 (n=51, Christensen et al) cohorts. The barplot shows the distribution of tumors according to pathological response (R) or absence of response (NR) in patients belonging to each consensus class. Patient’s response was defined by pathological downstaging to T1 or below. **(B)** Overall survival and response to PD-L1 blockade (Atezolizumab) in patients from the IMvigor210 trial (Mariathasan et al., 2018). We used our single-sample classifier to assign consensus classes to all IMvigor210 tumor samples profiled with RNA-seq. The barplot shows the distribution of tumors according to complete pathological response (CR), partial response (PR), stable disease (SD) or progressive disease (PD) in each consensus class. Consensus classes significantly associated with response to Atezolizumab (Fisher *P*<0.05), i.e. complete (CR) or partial responders (PR), are indicated by a black asterisk.

## Discussion

While precision genomic medicine promises to transform clinical practice, the diversity of published MIBC classifications has delayed transferring subtypes into both clinical trials and the standard management of bladder cancer patients. One approach to generating a standardized MIBC molecular classification would be to merge all available datasets and produce a new classification system. A concern for such an approach is that a range of methodologies could be applied. In the present study we chose a second approach, and generated a stable, unified classification system from the existing well-documented molecular subtypes. We analyzed the relationships among six published classification systems, based on 1750 MIBC transcriptomic profiles. We identified six consensus MIBC molecular classes that reconcile all six classification schemes: Basal/Squamous (Ba/Sq), Luminal Papillary (LumP), Luminal Unstable (LumU), Stroma-rich, Luminal Non-Specified (LumNS), and Neuroendocrine-like (NE-like). Each consensus class has distinct differentiation patterns, oncogenic mechanisms, tumor microenvironments, and histological and clinical associations (Figure 8). As well, we provide the community with an R-based, single-sample classifier that assigns a consensus class label to a tumor sample’s transcriptome.

**Figure 8:**
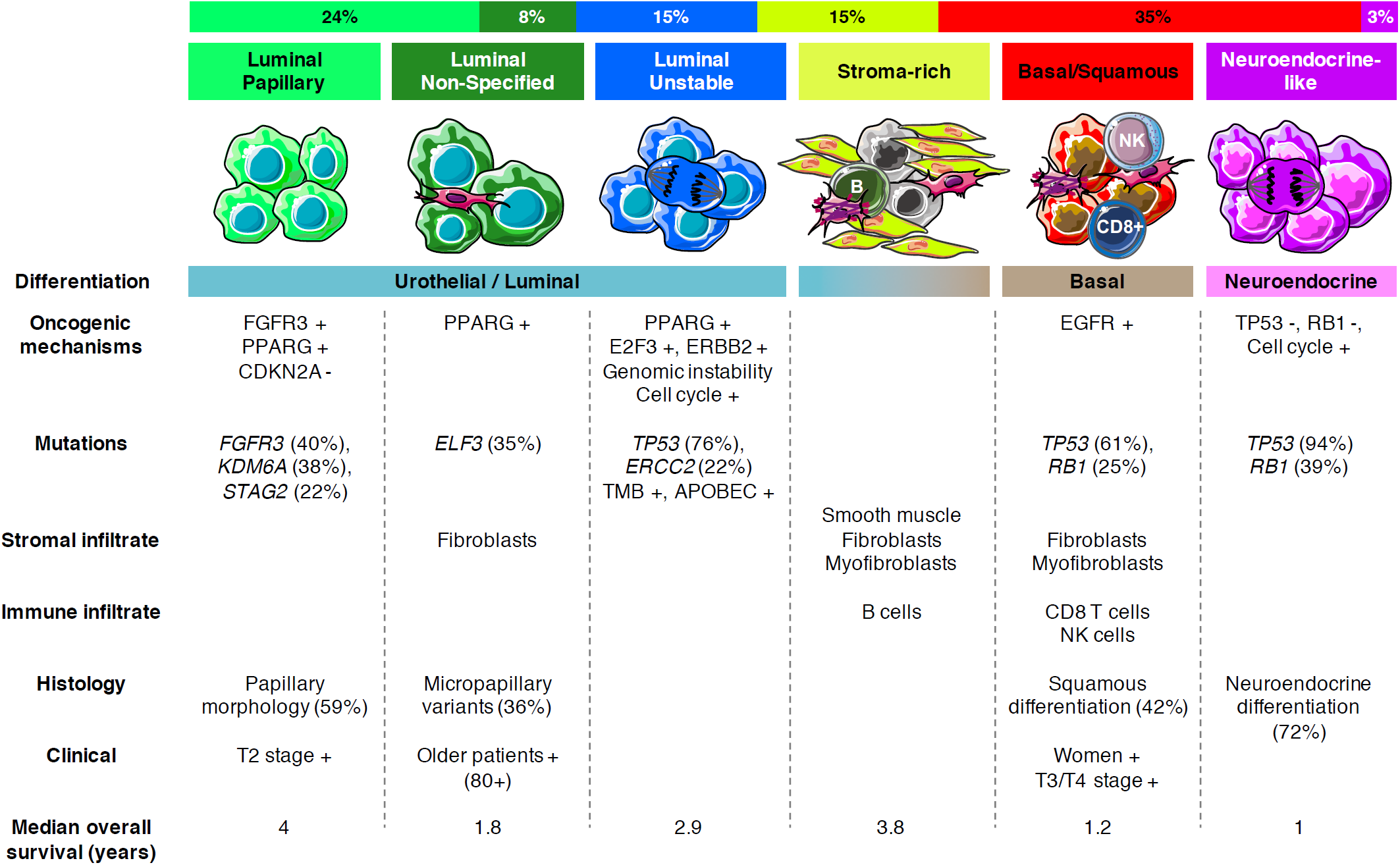
Summary of the main characteristics of the consensus classes. Top to bottom: Proportion of consensus classes in the n=1750 tumor samples. Consensus class names. Graphical display of tumor cells and their microenvironment (immune cells, fibroblasts, and smooth muscle cells). Differentiation-based color scale showing the features associated with consensus classes, including a Luminal-to-basal gradient and neuroendocrine differentiation. Table displaying the dominant characteristics: oncogenic mechanisms, mutations, stromal infiltrate, immune infiltrate, histology, clinical characteristics, and median overall survival.

The proposed consensus classification is consistent with MIBC differentiation-based stratification, revealing tumor classes that are primarily characterized by urothelial differentiation (Luminal classes), basal/squamous differentiation (Ba/Sq) and neuroendocrine differentiation (NE-like). Additional features, including genomic alterations, and pathological or clinical characteristics, are strongly associated with one or several classes (Figure 8).

LumP tumors are mainly characterized by strong transcriptional activation of *FGFR3*, involving a genetic mechanism (mutation, fusion, amplification) in more than 50% of LumP samples. Papillary morphology is more frequently observed in this consensus class. LumP tumors strongly express transcriptomic markers of the Ta pathway (Biton et al., 2014), and are consistently associated with the best prognosis compared to the other consensus classes. These data suggest that LumP tumors result from the progression of papillary Ta/T1 NMIBC.

The LumNS class includes a relatively small number of tumors, precluding using a more precise description. Nevertheless, our results point to interesting associations, such as an enrichment in *ELF3* mutations (35%) and an association with micropapillary morphology (36%), as well as a weak association with CIS (80%, n=4/5). The LumNS tumors are the only luminal tumors expressing stromal and immune signals. Their associated prognosis is the worst of the three luminal classes.

LumU tumors display typical features of genomic instability such as a higher tumor mutation burden, including more APOBEC-induced mutations, and more copy number alterations. The “Unstable” descriptor for this class refers to the Genomic Unstable tumors from the Lund classification, which are all included within this class. These tumors are particularly enriched in *TP53* and *ERCC2* mutations. LumU tumors are associated with high cell cycle activity, frequent E2F3/SOX4 amplifications, and *ERBB2* activation through mutations or amplification (63%).

Stroma-rich tumors are mainly characterized by high expression of non-tumor cell markers. Smooth muscle cells dominate the infiltration signals associated with these tumors, but endothelial cells and B lymphocytes are also overrepresented. As assessed by a urothelial differentiation signature (Eriksson et al., 2015) and by differentiation-based classification systems, this class contains both luminal and non-luminal tumors (Figure S7A). However, patients with luminal and non-luminal Stroma-rich tumors have very similar survival (Figure S7B). This suggests that although this subgroup is heterogeneous with regards to tumor cell phenotype, the presence of stroma could be the main parameter that drives its clinical features, given the current treatments.

Ba/Sq tumors have highest expression of basal differentiation markers and are strongly associated with squamous differentiation. Although the Ba/Sq tumors are characterized by high *KRT14, KRT5*/*6* and lack of *GATA3, FOXA1*, and *PPARG* expression, downregulation of *PPARG* and high expression of *KRT14* are not observed in normal basal cells (Fishwick et al., 2017). In this regard the Ba/Sq class is more similar to squamous urothelial metaplasia, consistent which enrichment in the squamous-cell carcinoma-associated C27 pan-cancer iCluster. Ba/Sq tumors express strong fibroblast and myofibroblast infiltration signals, as well as immune infiltration signals from cytotoxic T cells and NK cells. *EGFR* and *STAT3* activation are specific to this class.

The NE-like class includes virtually all (81%) of tumors with histological neuroendocrine differentiation. These tumors show high cell cycle activity, and all have both *TP53* and *RB1* genes inactivated by mutations or deletions. They have the worst prognosis of all MIBC consensus classes.

We generated the MIBC consensus classification following a procedure similar to that used to identify consensus subtypes in colorectal cancer (Guinney et al., 2015). The four Consensus Molecular Subtypes (CMS) identified in that work have helped frame the development of colorectal cancer precision medicine (Dienstmann et al., 2017) and are now being evaluated in clinical trials (Lenz et al., 2017; Mooi et al., 2018; Stintzing et al., 2017). Given the diverse nature of the six input classification systems that we used to define a consensus on MIBC molecular classes (distinct classification methods, strongly varying numbers of classes), we anticipate that the resulting consensus classification captures most of the molecular heterogeneity described and that it is currently a robust and useful solution for MIBC molecular classification.

Some bladder tumors show histological and molecular intra-tumor heterogeneity (Thomsen et al., 2017; Warrick et al., 2018). Our consensus subtyping system addresses inter-tumor heterogeneity, and focuses on defining the main molecular subtypes in MIBC. Our transcriptomic classifier will classify tumors according to the dominant class within the tumor sample analyzed. We recognize that tumor samples may contain multiple subtypes, and we address how such mixtures are likely to interfere with our single-sample classifier by having the classifier report not simply a class label, but also correlation values with all consensus classes. Further studies are required to assess the importance of intra-tumor heterogeneity in prognosis and response to treatment.

The consensus classification suggests possible therapeutic implications. Both the high rate of *FGFR3* mutations and translocations in LumP tumors, and the FGFR3 activation signature associated with these tumors, suggest that LumP tumors may respond to FGFR inhibitors, irrespective of the *FGFR3*’s mutation or translocation status. Novel fibroblast growth factor receptor inhibitors have been reported to clinically benefit MIBC patients that harbor mutations or translocations (about 20% of MIBC patients) and/or overexpression (about 40% of MIBC patients) of the tyrosine kinase receptor *FGFR3* (Nogova et al., 2017; Pal et al., 2018; Schuler et al., 2017; Tabernero et al., 2015).

There is increasing interest in targeting the tumor microenvironment, including the use of immunotherapy strategies. In the US and most of Europe, PD1 and PD-L1 immune checkpoint inhibition is becoming part of the standard of care for patients with locally advanced or metastatic urothelial cancer who relapse after cisplatin-based chemotherapy or are considered cisplatin-ineligible, with a 20% objective response rate. A phase 3 clinical trial has demonstrated the efficacy of targeting tumor vasculature in MIBC using an anti-VEGFR2 inhibitor (Petrylak et al., 2017). The different stromal components associated with the consensus classes, identified by transcriptomic signatures and revealed by our analysis of the IMvigor210 data, suggest that this consensus classification should be considered for further clinical studies involving immunotherapy or anti-angiogenic therapy.

Similarities between MIBC consensus classes and other cancer molecular subtypes may also be considered for future basket trials. We showed that such similarities are observed, for instance, between Ba/Sq MIBC tumors and squamous cell carcinomas arising in the head and neck, lung, and cervix, which were placed together in the C27 PanCanAtlas TCGA cluster. LumU tumors and other *ERBB2*-amplified tumors in breast and stomach cancers were also grouped together in the C2 TCGA PanCanAtlas cluster. More generally, bladder cancer and breast cancer luminal tumors share molecular similarities (Damrauer et al., 2014; Eriksson et al., 2015). Indeed, in both cancers the luminal subtypes rely on GATA3 and FOXA1, two transcription factors that are necessary for luminal differentiation, and on a nuclear receptor: the estrogen receptor (ESR1) in breast cancer and PPARG in bladder cancer (Biton et al., 2014). Intriguingly, in both cancers there is evidence that the nuclear receptor is involved in differentiation while also having pro-tumorigenic effects. Such comparisons across tumor types may help transfer treatment information from tumors bearing similar characteristics into bladder cancer.

We acknowledge limitations in the work reported here. Expression data were obtained with different methods and platforms. However, the approach applied to arrive at the consensus subtypes limits the possible biases caused by heterogenous data sources. Clinical data, on the other hand, are in most cases retrospectively collected, influenced by factors like patient selection, sample quality bias, missing data, and non-centralized pathology review, hard to correct for. Furthermore, the main outcome measure was overall survival, and for most patients we lacked detailed information on all treatments received during the disease course. Validating our findings, and determining whether subtype classification is an independent predictor of response or prognosticator of outcome, would require prospective studies in which the proposed classes are identified for patients who receive standardized treatments.

We emphasize that we report biological rather than clinical classes. We offer the mRNA classifier as a research tool for the retrospective and prospective exploratory work required to refine how such classes can best be used clinically. The consensus presented here provides a common foundation for the molecular classification of MIBC. Future sub-stratifications may allow defining a system that is more predictive of response to treatments; in such work, the clinical/strategic issue will be to decide the subtype granularity or resolution (Aine et al., 2015) that is appropriate for a specific problem. We expect this consensus classification to help the development of MIBC precision medicine by providing a robust framework to connect clinical findings to molecular contexts, and to identify clinically relevant biomarkers for patient management.

## Acknowledgements

This work is part of the French national program Cartes d’Identité des Tumeurs ^®^ (CIT) (http://cit.ligue-cancer.net). The results presented in this study are partly based upon data generated by The Cancer Genome Atlas (TCGA) pilot project established by the NCI and NHGRI. Information about TCGA and the investigators and institutions that constitute the TCGA research network can be found at http://cancergenome.nih.gov/. We are grateful for the patients who participated in this study. The results presented in this study are partly based on datasets financed by grants from the Swedish Cancer Society (2017/278) and Lund Medical Faculty (ALF).

## Author contributions

Conceptualization, all authors, initiated by F.X.R.; Methodology, A.K., A.dR., Y.A., G.S., K.A.H., K.S.C., W.Y.K., D.J.M., L.D., M.H., S.P.L., F.X.R., and F.R.; Software, A.K., A.dR., G.S., Q.M., J.K., W.C., and K.A.H.; Formal Analysis, A.K., A.dR., K.A.H., C.S.G, and M.A.A.C.; Investigation, A.K., A.dR., Y.A., and J.F.; Resources, Y.A., R.S., J.F., P.C.B., and L.D.; Writing - Review & Editing, all authors; Writing - Original Draft, A.K., A.dR., Y.A., G.S., A.G.R, L.D., M.H. and F.R.; Visualization, A.K., Y.A., G.S., and A.G.R.; Supervision, F.X.R. and F.R.

## Declaration of Interests

R.S. and P.C.B. share a patent with GenomeDX.

P.C.B. has been provided in-kind research funding by GenomeDX and is an advisory board member of AbbVie, Asieris, AstraZeneca, Astellas, Bayer, Biosyent, BMS, Janssen, Lilly, Merck, Roche, Sanofi, and Urogen and has participated in clinical trials with Genentech, Janssen, Ferring, Astellas, Sitka, and MDx Health.

J.B. has received research funding from Takeda, Pfizer, Novartis, and Sanofi-Aventis and has an advisory role for Genetech, MSD, Pfizer, GSK, BMS, Pierre Fabre, Sanofi-Aventis, Astellas, OncoGenex, and Janssen.

D.J.M. has received research funding from AstraZeneca and is a consultant for Rainier Pharmaceuticals.

S.P.L. is a consultant for UroGen and Vaxiion and has received support for a clinical trial from Endo, FKD, Urogen, and Viventia and is an advisory board member of Ferring, miR Scientific, QED Therapeutics, and UroGen.

## Methods

### Subtyping of MIBC samples according to published MIBC molecular classifications

Six methodologies for the molecular classification of MIBC samples using mRNA expression data had been previously developed and published among the expert teams contributing to this work. The six classification systems were built, as follows :

#### Baylor (Tumor differentiation)

Mo *et al* (Mo et al., 2018) developed a 18-gene tumor differentiation signature that molecularly define urothelial differentiation. This signature was used on 408 TCGA RNA-seq profiles to stratify MIBC patients into two groups, namely basal and differentiated. This classification is associated with distinct clinical outcomes, highlighting a correlation between MIBC tumor differentiation and its impact on patient’s survival. To classify a MIBC dataset of mRNA expression, they developed a classifier that performs a hierarchical clustering of sample profiles for the 18 genes, using complete linkage method and Pearson’s distance.

#### UNC

Damrauer *et al* (Damrauer et al., 2014) performed a consensus clustering on four aggregated gene expression array datasets, totaling 262 high-grade MIBC tumors. This analysis revealed two major clusters, termed luminal-like and basal-like, based on similarities with breast cancer subtypes. They developed a transcriptomic classifier using 47 genes (“BASE47”) to classify any transcriptomic profile of MIBC tumor. Into one of these two subtypes. The classifier was built using pamr R package.

#### CIT-Curie

Rebouissou *et al* (Rebouissou et al., 2014) performed a consensus hierarchical clustering in nine independent datasets including 370 gene expression microarray profiles of MIBC tumors. They measured the similarities between the mean expression profiles of the clusters identified in each dataset and defined seven meta-clusters (MC) which were represented in at least three datasets. The meta-cluster MC7 was further characterized as a molecular subtype of “Basal-like” MIBC tumors, activating EGFR oncogenic pathway. They built a transcriptomic classifier using nearest centroid method and Pearson’s correlation.

#### MDA

Choi *et al* (Choi et al., 2014) identified three subtypes through hierarchical clustering of a gene expression array dataset of 73 MIBC tumors. These subtypes were named basal, luminal and p53-like relatively to the transcriptomic markers and signatures expressed within each cluster. Basal and luminal subtypes shared similar molecular features with breast cancer subtypes, and p53-like tumors were shown to be resistant to chemotherapy. They built a transcriptomic classifier using nearest centroid method and Euclidean distance.

#### Lund

Marzouka *et al* (Marzouka et al., 2018) generated gene expression array data from 307 tumors and subdivided the cohort into six groups using stepwise hierarchical clustering on global gene expression. They further sub-stratified these groups into ten levels (‘tumor cell phenotypes”) using immunohistochemistry (IHC) data. By doing so, they show that their “Infiltrated group” can be assigned to a dominant tumor cell phenotype. The 10 resulting subtypes include urothelial-like subtypes named UroA-Prog, UroB, UroC, Uro-Inf (Infiltrated), GU (Genomically Unstable) and GU-Inf; and non-urothelial-like subtypes named Mes-like (Mesenchymal), Ba/Sq (Basal/Squamous), Ba/Sq-Inf, and Sc/NE-like (Small cell Neuroendocrine). The team built a transcriptomic based-classifier to assign these subtypes to any MIBC mRNA profile, using nearest centroid method and Pearson’s correlation.

#### TCGA

Robertson *et al* (Robertson et al., 2017) performed a NMF (Non-negative Matrix Factorization) consensus hierarchical clustering of RNA-seq profiles from 408 MIBC tumors compiled by TCGA and identified five expression subtypes. They further characterized these subtypes with extensive multi-omics data. They identified 3 luminal subtypes (named Luminal-papillary, Luminal-Infiltrated and Luminal), one “Basal/Squamous” subtype, and one “Neuronal” subtype. A transcriptomic classifier was built for the purpose of this consensus work and was validated by TCGA members. The classifier uses nearest centroid method and Pearson’s correlation.

Transcriptomic classifiers for Baylor, UNC, MDA, CIT-Curie, Lund, and TCGA classification systems were provided and/or validated by the respective teams. All classifiers were merged into an R package which is freely available at https://github.com/cit-bioinfo/BLCAsubtyping.

We used these classifiers on 18 MIBC mRNA datasets (*N* = 1750 samples) profiled on ten different gene expression platforms (Table S1), and assigned each sample to a subtype in each of the six classification systems (Figure S1). 16 datasets were retrieved from public repositories, and two unpublished datasets were shared by L.D. The normalization method applied on each dataset is detailed in Table S1. The six classifiers were applied on each dataset independently.

### Network construction and identification of consensus classes

Classification results from the six classifiers were merged for all 18 datasets, and transformed into a binary matrix D of 1750 samples (rows) × 29 classes (columns), where D(s, c) is set to 1 if sample s belongs to class c and 0 otherwise. Each row associated with a given sample contains exactly six 1’s, reflecting the six class labels predicted by the six classification systems. For each pair of classes, we computed Cohen’s Kappa scores to evaluate their agreement, i.e. to give a similarity measure between the corresponding pair of binary columns from the matrix. We built a weighted network, with 29 nodes encoding input molecular subtypes, and weighted edges encoding Cohen’s Kappa scores. If two subtypes were related by a Cohen’s Kappa score < 0, no edge was built between them, for negative values mean a complete absence of agreement between the two subtypes assignments. To quantify the statistical significance of the remaining edges, we performed hypergeometric tests for overrepresentation of samples classified to one subtype in another. The resulting p-values were adjusted for multiple hypothesis testing using the Benjamini–Hochberg (BH) method, and only edges corresponding to *P* < 0.001 were kept to build the network represented in Figure 2A. We used MCL(Van Dongen, 2008) (Markov cluster algorithm) to partition this network, as described previously (Guinney et al., 2015).

Clustering results were evaluated for MCL inflation factor I ranging from 3 to 15, with 0.3 increments, and using 500 resampling iterations. Each iteration consisted of randomly selecting 80% of samples before constructing the network and running MCL for a given inflation factor. For each inflation factor, we calculated a consensus clustering matrix, defined by the frequency that each pair of input subtypes is partitioned into the same cluster over all iterations. To evaluate clustering robustness, we computed a mean weighted silhouette width (MWSW) for each clustering result as previously described (Guinney et al., 2015). The weighted silhouette width extends the silhouette width by giving more weight to subtypes that are more stable within their assigned clusters. Here, we computed a stability score for each input subtype, defined as the average frequency over all iterations that its within-cluster associations with other subtypes is the same as predicted by MCL on the network generated with all samples. We then used these stability scores as subtypes’ weights to compute weighted silhouette width, and considered the mean over all subtypes as a measure of clustering robustness (Figure S2A). Clustering generated four-to six-cluster solutions, all of them yielding a mean silhouette width > 0.95 for at least one inflation factor value (Figure S2B). The K=4 solution was very robust (MWSW = 0.99) but poorly informative, revealing one cluster of basal subtypes, one cluster of luminal subtypes, one cluster of infiltrated classes, and one cluster of neuroendocrine-associated subtypes. The K=5 solution isolated an additional cluster containing only two subtypes (CIT MC7 and TCGA Luminal subtypes), which was not enough to clearly define a consensus class. The K=6 solution generated robust clusters (MWSW = 0.95), all containing a minimum of 3 subtypes). Heatmaps of consensus matrices for the three solutions illustrate the robustness of the clusters (Figure S2B). The 6-cluster solution was the optimal solution, yielding informative and robust clusters that were all supported by at least 3 input classification schemes.

### Identification of a core set of consensus samples

For each MIBC sample we performed a hypergeometric test for overrepresentation of the sample’s assigned input subtypes in the set of input subtypes associated with each consensus class. A sample was assigned to a consensus class if the corresponding overrepresentation test was significant (*P* < 0.001). Using this approach, we identified a core set of 1084 “consensus samples” to be highly representative of one of the 6 consensus classes. We used these consensus samples to build and validate a single-sample mRNA classifier for the consensus classes, then used this classifier to assign consensus labels to all 1750 MIBC samples.

### Single-sample transcriptomic classifier construction

We performed feature selection using a training subset of consensus samples from Sjödahl2017 (n=129) and TCGA (n=274) mRNA datasets, both of these sample sets including at least three consensus samples for each consensus class. In each dataset we performed LIMMA moderated t-tests (limma_3.39.1 R package) for each consensus class relative to the others and computed the AUC associated with each gene for the prediction of each class. We summarized the results for each gene common to both datasets (n=17381), using Stouffer’s method to aggregate p-values, and computing a mean fold-change for each class comparison. For each class, we selected the genes with Stouffer *P*<0.05 and AUC>0.6 in at least one of the two datasets, and ordered them according to their mean fold-change. We used these ordered gene lists to generate several lists of varying sizes, by selecting the N top upregulated genes and the N top down-regulated genes in each consensus class, with N varying from 10 to 125. A Pearson nearest-centroid classifier was built on the 129 Sjödahl2017 core samples for each of these gene lists, and its mean balanced accuracy was tested on the independent 681 consensus samples that had not been used for feature selection. The gene list that optimized mean balanced accuracy (97.23%, Figure S3) comprised 857 unique genes, and was used to build the final classifier. Six centroids corresponding to the six consensus classes (i.e. the mean mRNA profile of the 857 genes over each consensus class) were computed on the 129 consensus samples from Sjödahl2017 dataset. To classify the 1750 samples into one of the consensus classes, a Pearson correlation was computed between each sample and each centroid. Each sample was then assigned the consensus class whose centroid was the most correlated with the sample profile. If the maximal correlation for a given sample was less than 0.2, no consensus class label was assigned. This Pearson-based approach does not require to add a pre-processing step to the usual batch normalization of gene expression data, as long as the data are log-transformed, and can therefore be used in a single-sample setting. As shown in Figure S3C, the classifier accuracy was similar when using Affymetrix, Illumina, or RNA-seq data. The classifier is publicly available as an R package at https://github.com/cit-bioinfo/consensusMIBC.

### Comparison with TCGA pan-cancer classifications

The consensus bladder cancer classification schemed was compared to the TCGA’s PanCancerAtlas pan-cancer subtypes (Hoadley et al., 2018). We visualized the overlap of classification schemes by calculating the percentage within each MIBC consensus class across the TCGA Pan-Cancer Atlas iCluster classification. We then normalized each row (consensus class) by setting the sum of squares equal to 1. We clustered these data using 1-pearson correlation and used a heatmap for visualization. To evaluate the significance of the enrichment of consensus classes with certain pan-cancer classifications, we calculated the Chi-Square or Fisher’s Exact test p-value from a 2×2 contingency table for the given two classifications of interest. To account for multiple testing, we calculated the Bonferroni p-value threshold for 441 pairwise comparisons to be *P*<0.00011.

### Extraction of bladder cancer gene signatures from Biton *et al*

In their study, Biton *et al* identified and characterized several major bladder cancer signals by an independent component analysis of bladder cancer transcriptome data (Biton et al., 2014). We used ten of these independent components to extract gene sets associated with the Ta pathway (CIT-13), basal differentiation (CIT-6), cell cycle (CIT-7), urothelial differentiation (CIT-9), smooth muscle (CIT-3), lymphocytes B&T (CIT-8), myofibroblasts (CIT-12), interferon response (CIT-5), neuroendocrine differentiation (CIT-18), and mitochondria (CIT-4). We retrieved the sample contribution vectors associated with each of these components and correlated these values to each gene of the CIT mRNA dataset. Genes that had a Pearson correlation greater than 0.6 (or less than −0.6, depending on the direction of the component association with the biological signal) were selected as representative gene sets for the biological signals associated to the component. The resulting gene sets are given in Table S3.

### Significance analysis for gene sets differential expression

For each mRNA dataset, we used the R package GSA (Efron and Tibshirani, 2007) (1.03.1) to compare the differential expression of gene sets (Table S3 and Table S7) in each consensus classes relatively to the others. The method returns 2 p-values by gene set, assessing the significance of lower (respectively higher) expression of the gene set between two conditions. We used Stouffer’s method to summarize these p-values across the 18 datasets analyzed, yielding 2 aggregated p-values (significance assessment of higher and lower expression) for each gene set.

### Computation of regulon activity scores for 23 regulators

A transcriptional regulatory network for 23 regulators reported as associated with bladder cancer was reconstructed from the TCGA (n=404) MIBC RNA-seq data (Robertson et al., 2017) using the RTN R package (2.6.0). This regulatory network reconstruction was provided as an RTN TNI-class object, and used to calculate regulon activity scores for 18 cohorts, individually. In each sample in each cohort, for each regulon we used RTN’s tni.gsea2 function to calculate two-tailed GSEA tests (Castro et al., 2016). This generated regulon activity profiles (RAPs) for each cohort; such a profile shows regulon activities of samples, relative to other samples in the same cohort. Regulons were also assigned discrete status as ‘activated’, ‘neutral’ and ‘inactivated’ in each sample based on their activity.

### Statistical analyses

We measured association between consensus classes and categorical variables by Fisher’s exact or Chi-square tests. We evaluated differences of continuous variables distributions between consensus classes by Kruskal-Wallis tests, ANOVA or LIMMA moderated t-tests (limma_3.39.1 R package).

We built multivariate Cox models integrating consensus classes and clinical risk factors, stratified on cohort of patients (separate baseline hazard functions were fit for each strata). We used Wald tests to assess survival differences associated with different levels of a given factor included in the Cox models. For each factor level, we computed Hazard Ratios (HR) and 95% Confidence Intervals (CI). We constructed Kaplan-Meier curves to visualize overall survival stratified by consensus class and used log-rank tests to compare the survival of corresponding patient groups.

All statistical and bioinformatics analyses were performed with R software environment (version 3.5.1).

### Code availability

Subtyping algorithms for the 6 input classification systems were implemented within R software environment (version 3.5.1) and have been wrapped into an R package freely available at https://github.com/cit-bioinfo/BLCAsubtyping.

MCL algorithm (version mcl-14-137) was used for the discovery of consensus classes and is available at https://micans.org/mcl/.

The transcriptomic classifier to assign consensus classes was developed within R software environment (version 3.5.1) and is freely available at https://github.com/cit-bioinfo/consensusMIBC.

GISTIC 2 (release 2.0.23) was used to analyze segmented copy number data and find recurrent genomic alterations.

All other analyses were performed using R software environment (version 3.5.1) and freely available packages from either The Comprehensive R Archive Network (CRAN, https://cran.r-project.org/web/packages/available_packages_by_name.html) or Bioconductor repository (https://bioconductor.org/packages/release/bioc/). Non basic R packages used for the analyses have been all properly mentioned throughout the appropriate method sections.

### Data availability

The 16 public datasets analyzed to derive the consensus classification are available under accession names provided in Table S1. Aarhus datasets 1 and 2 are available from L.D. on reasonable request. Aarhus.3 dataset is to be published in Journal of Clinical Oncology (Christensen et al). MDA MVAC and DDMVAC datasets were downloaded from GEO repository through accession names GSE70691 and GSE69795. IMvigor210 dataset was downloaded from the data folder of the IMvigor210CoreBiologies R package (version 0.1.13, http://research-pub.gene.com/IMvigor210CoreBiologies/packageVersions/).

## Supplemental Information titles and legends

**Figure S1. Related to Figure 1; Subtyping results using the six input classification systems (Baylor, CIT-Curie, UNC, Lund, MDA, and TCGA).** The six classifiers are described in Methods and were run independently on each of the 18 datasets described in Table S1. The distributions of subtyping results by dataset are represented by the barplots (one barplot by classification system).

**Figure S2. Related to Figure 1; MCL clustering results**. (**A**) Clustering performance: mean weighted silhouette width of the network clustering, as a function of inflation factor I. The number plotted above each point gives the number of resulting classes. All clustering results had a very high silhouette width (>0.88). We selected the inflation factor that maximized the mean weighted silhouette width among the 6-classes solutions, which corresponds to a local maximum of the plotted function (I = 11.4, red dot). (**B**) Consensus clustering matrices for the best 4-class (I = 3), 5-class (I = 4.8) and 6-class (I = 11.4) solutions (related to bold dots in (**A**)).

**Figure S3. Related to Figure 2; Classifier performance metrics.** A single-sample classifier was built on a subset of 403 core consensus samples (129 samples from the Sjodahl2017 dataset and 274 samples from the TCGA dataset), and was tested on the remaining 681 core consensus samples from the 16 other datasets. Balanced accuracy, specificity, and sensitivity were measured regarding prediction of the 6 consensus classes (**A**) on validation samples only (**B**) and on the whole set of core consensus samples. (**C**) Performance metrics for Affymetrix, Illumina, and RNA-seq profiling.

**Figure S4. Related to Figure 2; Distribution of consensus class predictions across initial subtypes, across each dataset, and relationship with TCGA pan-cancer classes. (A)** Consensus class predictions within each subtype of the six original classification systems. **(B)** Consensus class predictions within each of the 18 datasets described in Table S1. **(C)** Enrichment of MIBC tumors from consensus classes in each of the TCGA PanCanAtlas integrative clusters (Hoadley et al., 2018). Enrichment was evaluated using 406 MIBC tumors from the TCGA BLCA dataset. We excluded the 18 of 33 PanCanAtlas iCluster groups which did not include any bladder tumors.

**Figure S5. Related to Figure 4; Analysis of copy number and exome data, and associations with consensus classes. (A)** Sample tumor purity across consensus classes, as estimated by ABSOLUTE on TCGA copy number data. ABSOLUTE estimates were available for 397 TCGA MIBC samples through Robertson et al publication data. (**B**) Distribution of Somatic Copy Number Alteration (SCNA) counts across consensus classes. SCNA counts are defined as the number of genes with copy number changes, as estimated by GISTIC2 over 600 MIBC CNV profiles from CIT (n=87), Iyer (n=58), Sjödahl2012 (n =29), Stransky (n=22) and TCGA (n=404) datasets. (**C**) Distribution of nonsynonymous somatic mutation events across consensus classes. The boxes in (a), (b), and (c) show the 25th percentile, 50th percentile, and 75th percentile of the data, with ‘‘whiskers’’ extending to the most extreme data point which is no more than 1.5 times the inter-quartile range. (**D**) Enrichment of APOBEC-induced mutations within consensus classes. The minimum estimate of the number of APOBEC-induced mutations was computed for 406 samples of TCGA MIBC cohort and discretized into categorical values: “No” : estimate = 0; “Low“: estimate ≤ median of non-zero values (median was 61.5); “High“: estimate > median of non-zero values. These values were retrieved from Robertson et al publication data.

**Figure S6. Related to Figure 5; The three histological features most associated with consensus classes**. Example micrographs of the three histological features that were most significantly associated with one of the consensus classes: **(A)** Squamous differentiation, **(B)** Micropapillary variants, and **(B)** Neuroendocrine differentiation. These pictures were obtained from TCGA tissue sample digital images (sample names are given below each picture) which are available through the Cancer Digital Slide Archive (cancer.digitalslidearchive.net). For each feature, a barplot shows the percentage of histologically reviewed tumors in each consensus class that present this histological characteristic. Fisher’s exact tests were performed to assess the strength of enrichment within each group; a p-value is given below each barplot for the most significant association (indicated with an asterisk). The total number of cases reviewed for each feature is given in parenthesis under each barplot.

**Figure S7. Related to Figure 6; Differentiation status (luminal vs non-luminal) of Stroma-rich tumors and associated overall survival. (A)** Consistency of three independent methods to assess the dominant differentiation status of 263 stroma-rich samples. The two top layers of the heatmap give the sample subtype (Basal or Luminal/Differentiated) according to UNC and Baylor (Tumor differentiation) classifications. The bottom gradient shows the mean expression of an urothelial differentiation signature (Eriksson et al.,2015; Table S3), scaled on each of the 18 datasets analyzed. (B) Overall survival of patients with stroma-rich tumors, stratified by tumor differentiation related subtypes, using either UNC classification or Baylor (Tumor differentiation) classification.

**Table S1. Related to Figure 1; Description of the 18 mRNA datasets and MIBC tumors included for the consensus discovery.** 16 datasets are publicly available using the information in “Data accession column”. The number of muscle-invasive samples used from each dataset is given in the “MIBC samples” column. Samples originating from post-treatment tumor samples were excluded. Available clinical characteristics associated to the included samples are summarized in the four last columns.

**Table S2. Related to Figure 2; Consensus classifier calls for the 1750 MIBC samples from the 18 datasets analyzed, and additional clinical, molecular, and histological data used in the study.**

**Table S3. Related to Figure 3; Description of the transcriptomic signatures summarized in Figure 3, and associated p-values.** For the gene sets analyses (Biton et al, Eriksson et al, Sjödahl et al signatures), we used Stouffer’s method to combine GSA p-values for lower expression (respectively higher expression) in each consensus class relatively to the others, after independent analyses of 18 datasets. For the computational tools MCP counter and ESTIMATE, we used Stouffer’s method to combine t-test p-values for lower scores (respectively higher scores) in each consensus class relatively to the others, after independent analyses of 18 datasets. For regulon status analysis, we used Fisher’s exact tests to assess over-enrichment of inactive (respectively active) regulon status in each consensus class relatively to the others.

**Table S4. Related to Figure 4; Association of the consensus molecular classification with exome data from TCGA (n=388).** We used processed exome TCGA data to analyze the mutation profiles relatively to each consensus class. Only genes with MutsigCV Q-values < 0.02 (computed by TCGA) are listed in the table. Chi-squared tests were performed for each gene to test if the gene mutations were differentially enriched within the consensus classes. For each class, Fisher’s exact tests were performed to measure specific enrichment in the class.

**Table S5. Related to Figure 4; Regions of amplification or deletion identified by GISTIC2 algorithm and significantly associated with one of the consensus classes.** We analyzed 600 segmented copy number profiles from 5 datasets (TCGA, CIT, Stransky, Sjödahl, Iyer). GISTIC2 was used to identify significantly altered regions within each consensus class (GISTIC residual q-value < 0.05) and Fisher exact tests were performed to test the specific association of the regions with each consensus class compared to the others. For each regions we computed the gene expression vs copy number (CN) correlation of all encompassed genes and marked as candidate genes those whose Pearson correlation was at least 0.4 in one of the 5 datasets analyzed. The best observed correlation among all genes within the region is reported here, alongside with the name of corresponding gene. For each consensus class, we highlighted in bold the region whose copy number aberrations were the most significantly associated with the class.

**Table S6. Related to Figure 6; Univariate and multivariate cox regression models.** Patients who were said to have received neoadjuvant chemotherapy were excluded from the survival analyses (a) Univariate Cox regression models were performed based on 873 MIBC patients with available clinical data. Factors significantly associated with survival have likelihood p-values highlighted in bold letters. Only clinical factors that were significant in the univariate Cox regressions were kept to build the multivariate Cox model in (b), which included 509 patients with complete clinical information. A multivariate Cox model was also built for each consensus class separately to assess the added prognostic value of each class subtyping as compared to the other clinical factors.

**Table S7. Related to Figure 6; Description and statistics associated with the gene signatures summarized in Figure 6c.** The Stouffer aggregated p-values combine GSA p-values for lower expression (respectively higher expression) obtained through the independent analyses performed on 18 datasets to test the differential expression of the listed gene sets in each consensus class relatively to the others. For the differential expression of individual genes (PD-1 and PD-L1), the Stouffer p-values combine one-sided t-tests p-values performed on each dataset.

**Supplementary Note. Related to Figure 2.**

## References

Aine, M., Eriksson, P., Liedberg, F., Höglund, M., and Sjödahl, G. (2015). On Molecular Classification of Bladder Cancer: Out of One, Many. Eur. Urol. 68, 921–923.

Ayers, M., Lunceford, J., Nebozhyn, M., Murphy, E., Loboda, A., Kaufman, D.R., Albright, A., Cheng, J.D., Kang, S.P., Shankaran, V., et al. (2017). IFN-γ-related mRNA profile predicts clinical response to PD-1 blockade. J. Clin. Invest. 127, 2930–2940.

Becht, E., Giraldo, N.A., Lacroix, L., Buttard, B., Elarouci, N., Petitprez, F., Selves, J., Laurent-Puig, P., Sautès-Fridman, C., Fridman, W.H., et al. (2016). Estimating the population abundance of tissue-infiltrating immune and stromal cell populations using gene expression. Genome Biol. 17, 218.

Biton, A., Bernard-Pierrot, I., Lou, Y., Krucker, C., Chapeaublanc, E., Rubio-Pérez, C., López-Bigas, N., Kamoun, A., Neuzillet, Y., Gestraud, P., et al. (2014). Independent component analysis uncovers the landscape of the bladder tumor transcriptome and reveals insights into luminal and basal subtypes. Cell Rep. 9, 1235–1245.

Blaveri, E., Simko, J.P., Korkola, J.E., Brewer, J.L., Baehner, F., Mehta, K., Devries, S., Koppie, T., Pejavar, S., Carroll, P., et al. (2005). Bladder cancer outcome and subtype classification by gene expression. Clin. Cancer Res. Off. J. Am. Assoc. Cancer Res. 11, 4044–4055.

Böck, M., Hinley, J., Schmitt, C., Wahlicht, T., Kramer, S., and Southgate, J. (2014). Identification of ELF3 as an early transcriptional regulator of human urothelium. Dev. Biol. 386, 321–330.

Cancer Genome Atlas Research Network (2014). Comprehensive molecular characterization of urothelial bladder carcinoma. Nature 507, 315–322.

Carter, S.L., Cibulskis, K., Helman, E., McKenna, A., Shen, H., Zack, T., Laird, P.W., Onofrio, R.C., Winckler, W., Weir, B.A., et al. (2012). Absolute quantification of somatic DNA alterations in human cancer. Nat. Biotechnol. 30, 413–421.

Castro, M.A.A., de Santiago, I., Campbell, T.M., Vaughn, C., Hickey, T.E., Ross, E., Tilley, W.D., Markowetz, F., Ponder, B.A.J., and Meyer, K.B. (2016). Regulators of genetic risk of breast cancer identified by integrative network analysis. Nat. Genet. 48, 12–21.

Choi, W., Porten, S., Kim, S., Willis, D., Plimack, E.R., Hoffman-Censits, J., Roth, B., Cheng, T., Tran, M., Lee, I.-L., et al. (2014). Identification of distinct basal and luminal subtypes of muscle-invasive bladder cancer with different sensitivities to frontline chemotherapy. Cancer Cell 25, 152–165.

Choi, W., Ochoa, A., McConkey, D.J., Aine, M., Höglund, M., Kim, W.Y., Real, F.X., Kiltie, A.E., Milsom, I., Dyrskjøt, L., et al. (2017). Genetic Alterations in the Molecular Subtypes of Bladder Cancer: Illustration in the Cancer Genome Atlas Dataset. Eur. Urol. 72, 354–365.

Damrauer, J.S., Hoadley, K.A., Chism, D.D., Fan, C., Tiganelli, C.J., Wobker, S.E., Yeh, J.J., Milowsky, M.I., Iyer, G., Parker, J.S., et al. (2014). Intrinsic subtypes of high-grade bladder cancer reflect the hallmarks of breast cancer biology. Proc. Natl. Acad. Sci. U. S. A. 111, 3110–3115.

Dienstmann, R., Vermeulen, L., Guinney, J., Kopetz, S., Tejpar, S., and Tabernero, J. (2017). Consensus molecular subtypes and the evolution of precision medicine in colorectal cancer. Nat. Rev. Cancer 17, 79–92.

Dyrskjøt, L., Thykjaer, T., Kruhøffer, M., Jensen, J.L., Marcussen, N., Hamilton-Dutoit, S., Wolf, H., and Orntoft, T.F. (2003). Identifying distinct classes of bladder carcinoma using microarrays. Nat. Genet. 33, 90–96.

Efron, B., and Tibshirani, R. (2007). On testing the significance of sets of genes. Ann. Appl. Stat. 1, 107–129.

Eriksson, P., Aine, M., Veerla, S., Liedberg, F., Sjödahl, G., and Höglund, M. (2015). Molecular subtypes of urothelial carcinoma are defined by specific gene regulatory systems. BMC Med. Genomics 8, 25.

Fishwick, C., Higgins, J., Percival-Alwyn, L., Hustler, A., Pearson, J., Bastkowski, S., Moxon, S., Swarbreck, D., Greenman, C.D., and Southgate, J. (2017). Heterarchy of transcription factors driving basal and luminal cell phenotypes in human urothelium. Cell Death Differ. 24, 809–818.

Guinney, J., Dienstmann, R., Wang, X., de Reyniès, A., Schlicker, A., Soneson, C., Marisa, L., Roepman, P., Nyamundanda, G., Angelino, P., et al. (2015). The consensus molecular subtypes of colorectal cancer. Nat. Med. 21, 1350–1356.

Hedegaard, J., Lamy, P., Nordentoft, I., Algaba, F., Høyer, S., Ulhøi, B.P., Vang, S., Reinert, T., Hermann, G.G., Mogensen, K., et al. (2016). Comprehensive Transcriptional Analysis of Early-Stage Urothelial Carcinoma. Cancer Cell 30, 27–42.

Hoadley, K.A., Yau, C., Hinoue, T., Wolf, D.M., Lazar, A.J., Drill, E., Shen, R., Taylor, A.M., Cherniack, A.D., Thorsson, V., et al. (2018). Cell-of-Origin Patterns Dominate the Molecular Classification of 10,000 Tumors from 33 Types of Cancer. Cell 173, 291–304.e6.

Horsman, M.R., and Overgaard, J. (2016). The impact of hypoxia and its modification of the outcome of radiotherapy. J. Radiat. Res. (Tokyo) 57, i90–i98.

Hurst, C.D., Alder, O., Platt, F.M., Droop, A., Stead, L.F., Burns, J.E., Burghel, G.J., Jain, S., Klimczak, L.J., Lindsay, H., et al. (2017). Genomic Subtypes of Non-invasive Bladder Cancer with Distinct Metabolic Profile and Female Gender Bias in KDM6A Mutation Frequency. Cancer Cell 32, 701–715.e7.

Kim, J., Kwiatkowski, D., McConkey, D.J., Meeks, J.J., Freeman, S.S., Bellmunt, J., Getz, G., and Lerner, S.P. (2019). The Cancer Genome Atlas Expression Subtypes Stratify Response to Checkpoint Inhibition in Advanced Urothelial Cancer and Identify a Subset of Patients with High Survival Probability. Eur. Urol.

Knowles, M.A., and Hurst, C.D. (2015). Molecular biology of bladder cancer: new insights into pathogenesis and clinical diversity. Nat. Rev. Cancer 15, 25–41.

Lenz, H.-J., Ou, F.-S., Venook, A.P., Hochster, H.S., Niedzwiecki, D., Goldberg, R.M., Mayer, R.J., Bertagnolli, M.M., Blanke, C.D., Zemla, T., et al. (2017). Impact of consensus molecular subtyping (CMS) on overall survival (OS) and progression free survival (PFS) in patients (pts) with metastatic colorectal cancer (mCRC): Analysis of CALGB/SWOG 80405 (Alliance). J. Clin. Oncol. 35, 3511–3511.

Lindgren, D., Frigyesi, A., Gudjonsson, S., Sjödahl, G., Hallden, C., Chebil, G., Veerla, S., Ryden, T., Månsson, W., Liedberg, F., et al. (2010). Combined gene expression and genomic profiling define two intrinsic molecular subtypes of urothelial carcinoma and gene signatures for molecular grading and outcome. Cancer Res. 70, 3463–3472.

Mariathasan, S., Turley, S.J., Nickles, D., Castiglioni, A., Yuen, K., Wang, Y., Iii, E.E.K., Koeppen, H., Astarita, J.L., Cubas, R., et al. (2018). TGFβ attenuates tumour response to PD- L1 blockade by contributing to exclusion of T cells. Nature 554, 544–548.

Marzouka, N., Eriksson, P., Rovira, C., Liedberg, F., Sjödahl, G., and Höglund, M. (2018). A validation and extended description of the Lund taxonomy for urothelial carcinoma using the TCGA cohort. Sci. Rep. 8, 3737.

Mermel, C.H., Schumacher, S.E., Hill, B., Meyerson, M.L., Beroukhim, R., and Getz, G. (2011). GISTIC2.0 facilitates sensitive and confident localization of the targets of focal somatic copy-number alteration in human cancers. Genome Biol. 12, R41.

Mo, Q., Nikolos, F., Chen, F., Tramel, Z., Lee, Y.-C., Hayashi, K., Xiao, J., Shen, J., and Chan, K.S. (2018). Prognostic Power of a Tumor Differentiation Gene Signature for Bladder Urothelial Carcinomas. J. Natl. Cancer Inst.

Mooi, J.K., Wirapati, P., Asher, R., Lee, C.K., Savas, P., Price, T.J., Townsend, A., Hardingham, J., Buchanan, D., Williams, D., et al. (2018). The prognostic impact of consensus molecular subtypes (CMS) and its predictive effects for bevacizumab benefit in metastatic colorectal cancer: molecular analysis of the AGITG MAX clinical trial. Ann. Oncol. Off. J. Eur. Soc. Med. Oncol. 29, 2240–2246.

Nogova, L., Sequist, L.V., Perez Garcia, J.M., Andre, F., Delord, J.-P., Hidalgo, M., Schellens, J.H.M., Cassier, P.A., Camidge, D.R., Schuler, M., et al. (2017). Evaluation of BGJ398, a Fibroblast Growth Factor Receptor 1-3 Kinase Inhibitor, in Patients With Advanced Solid Tumors Harboring Genetic Alterations in Fibroblast Growth Factor Receptors: Results of a Global Phase I, Dose-Escalation and Dose-Expansion Study. J. Clin. Oncol. Off. J. Am. Soc. Clin. Oncol. 35, 157–165.

Pal, S.K., Rosenberg, J.E., Hoffman-Censits, J.H., Berger, R., Quinn, D.I., Galsky, M.D., Wolf, J., Dittrich, C., Keam, B., Delord, J.-P., et al. (2018). Efficacy of BGJ398, a fibroblast growth factor receptor 1-3 inhibitor, in patients with previously treated advanced urothelial carcinoma with FGFR3 alterations. Cancer Discov. CD-18-0229.

Pawlik, T.M., and Keyomarsi, K. (2004). Role of cell cycle in mediating sensitivity to radiotherapy. Int. J. Radiat. Oncol. Biol. Phys. 59, 928–942.

Petrylak, D.P., Wit, R. de, Chi, K.N., Drakaki, A., Sternberg, C.N., Nishiyama, H., Castellano, D., Hussain, S., Fléchon, A., Bamias, A., et al. (2017). Ramucirumab plus docetaxel versus placebo plus docetaxel in patients with locally advanced or metastatic urothelial carcinoma after platinum-based therapy (RANGE): a randomised, double-blind, phase 3 trial. The Lancet 390, 2266–2277.

Rebouissou, S., Bernard-Pierrot, I., de Reyniès, A., Lepage, M.-L., Krucker, C., Chapeaublanc, E., Hérault, A., Kamoun, A., Caillault, A., Letouzé, E., et al. (2014). EGFR as a potential therapeutic target for a subset of muscle-invasive bladder cancers presenting a basal-like phenotype. Sci. Transl. Med. 6, 244ra91.

Robertson, A.G., Kim, J., Al-Ahmadie, H., Bellmunt, J., Guo, G., Cherniack, A.D., Hinoue, T., Laird, P.W., Hoadley, K.A., Akbani, R., et al. (2017). Comprehensive Molecular Characterization of Muscle-Invasive Bladder Cancer. Cell 171, 540–556.e25.

Rosenberg, J.E., Hoffman-Censits, J., Powles, T., Heijden, M.S. van der, Balar, A.V., Necchi, A., Dawson, N., O’Donnell, P.H., Balmanoukian, A., Loriot, Y., et al. (2016). Atezolizumab in patients with locally advanced and metastatic urothelial carcinoma who have progressed following treatment with platinum-based chemotherapy: a single-arm, multicentre, phase 2 trial. The Lancet 387, 1909–1920.

Schuler, M., Nogova, L., Heidenreich, A., Tai, D., Cassier, P., Richly, H., Cho, B.C., Sayehli, C.M., Navarro, A., Bender, S., et al. (2017). 859PAnti-tumor activity of the pan-FGFR inhibitor rogaratinib in patients with advanced urothelial carcinomas selected based on tumor FGFR mRNA expression levels. Ann. Oncol. 28.

Seiler, R., Ashab, H.A.D., Erho, N., Rhijn, B.W.G. van, Winters, B., Douglas, J., Kessel, K.E.V., Putte, E.E.F. van de, Sommerlad, M., Wang, N.Q., et al. (2017). Impact of Molecular Subtypes in Muscle-invasive Bladder Cancer on Predicting Response and Survival after Neoadjuvant Chemotherapy. Eur. Urol. 72, 544–554.

Sjödahl, G., Lauss, M., Lövgren, K., Chebil, G., Gudjonsson, S., Veerla, S., Patschan, O., Aine, M., Fernö, M., Ringnér, M., et al. (2012). A molecular taxonomy for urothelial carcinoma. Clin. Cancer Res. Off. J. Am. Assoc. Cancer Res. 18, 3377–3386.

Sjödahl, G., Eriksson, P., Liedberg, F., and Höglund, M. (2017). Molecular classification of urothelial carcinoma: global mRNA classification versus tumour-cell phenotype classification. J. Pathol. 242, 113–125.

Stintzing, S., Wirapati, P., Lenz, H.-J., Neureiter, D., Fischer von Weikersthal, L., Decker, T., Kiani, A., Vehling-Kaiser, U., Al-Batran, S.-E., Heintges, T., et al. (2017). Consensus molecular subgroups (CMS) of colorectal cancer (CRC) and first-line efficacy of FOLFIRI plus cetuximab or bevacizumab in the FIRE3 (AIO KRK-0306) trial. J. Clin. Oncol. 35, 3510–3510.

Tabernero, J., Bahleda, R., Dienstmann, R., Infante, J.R., Mita, A., Italiano, A., Calvo, E., Moreno, V., Adamo, B., Gazzah, A., et al. (2015). Phase I Dose-Escalation Study of JNJ- 42756493, an Oral Pan–Fibroblast Growth Factor Receptor Inhibitor, in Patients With Advanced Solid Tumors. J. Clin. Oncol. 33, 3401–3408.

Tan, T.Z., Rouanne, M., Tan, K.T., Huang, R.Y.-J., and Thiery, J.-P. (2019). Molecular Subtypes of Urothelial Bladder Cancer: Results from a Meta-cohort Analysis of 2411 Tumors. Eur. Urol. 75, 423–432.

Thomsen, M.B.H., Nordentoft, I., Lamy, P., Vang, S., Reinert, L., Mapendano, C.K., Høyer, S., Ørntoft, T.F., Jensen, J.B., and Dyrskjøt, L. (2017). Comprehensive multiregional analysis of molecular heterogeneity in bladder cancer. Sci. Rep. 7, 11702.

Van Dongen, S. (2008). Graph Clustering Via a Discrete Uncoupling Process. SIAM J. Matrix Anal. Appl. 30, 121–141.

Volkmer, J.-P., Sahoo, D., Chin, R.K., Ho, P.L., Tang, C., Kurtova, A.V., Willingham, S.B., Pazhanisamy, S.K., Contreras-Trujillo, H., Storm, T.A., et al. (2012). Three differentiation states risk-stratify bladder cancer into distinct subtypes. Proc. Natl. Acad. Sci. U. S. A. 109, 2078–2083.

Warrick, J.I., Sjödahl, G., Kaag, M., Raman, J.D., Merrill, S., Shuman, L., Chen, G., Walter, V., and DeGraff, D.J. (2018). Intratumoral Heterogeneity of Bladder Cancer by Molecular Subtypes and Histologic Variants. Eur. Urol. 0.

Yoshihara, K., Shahmoradgoli, M., Martínez, E., Vegesna, R., Kim, H., Torres-Garcia, W., Treviño, V., Shen, H., Laird, P.W., Levine, D.A., et al. (2013). Inferring tumour purity and stromal and immune cell admixture from expression data. Nat. Commun. 4, 2612.

