## Supplementary figures and images for "A consensus molecular classification of muscle-invasive bladder cancer"

### Figure S1

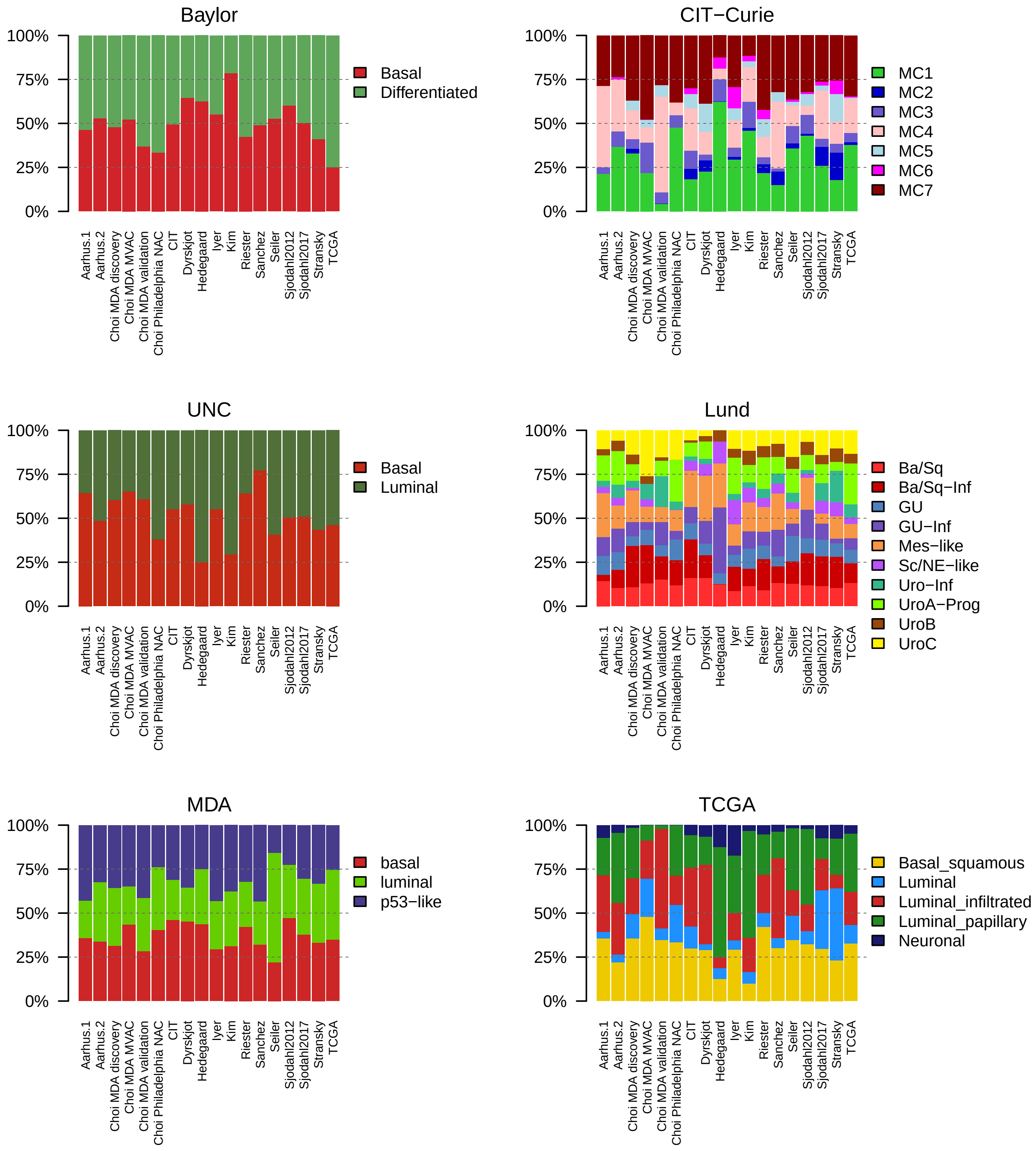

### Figure S2

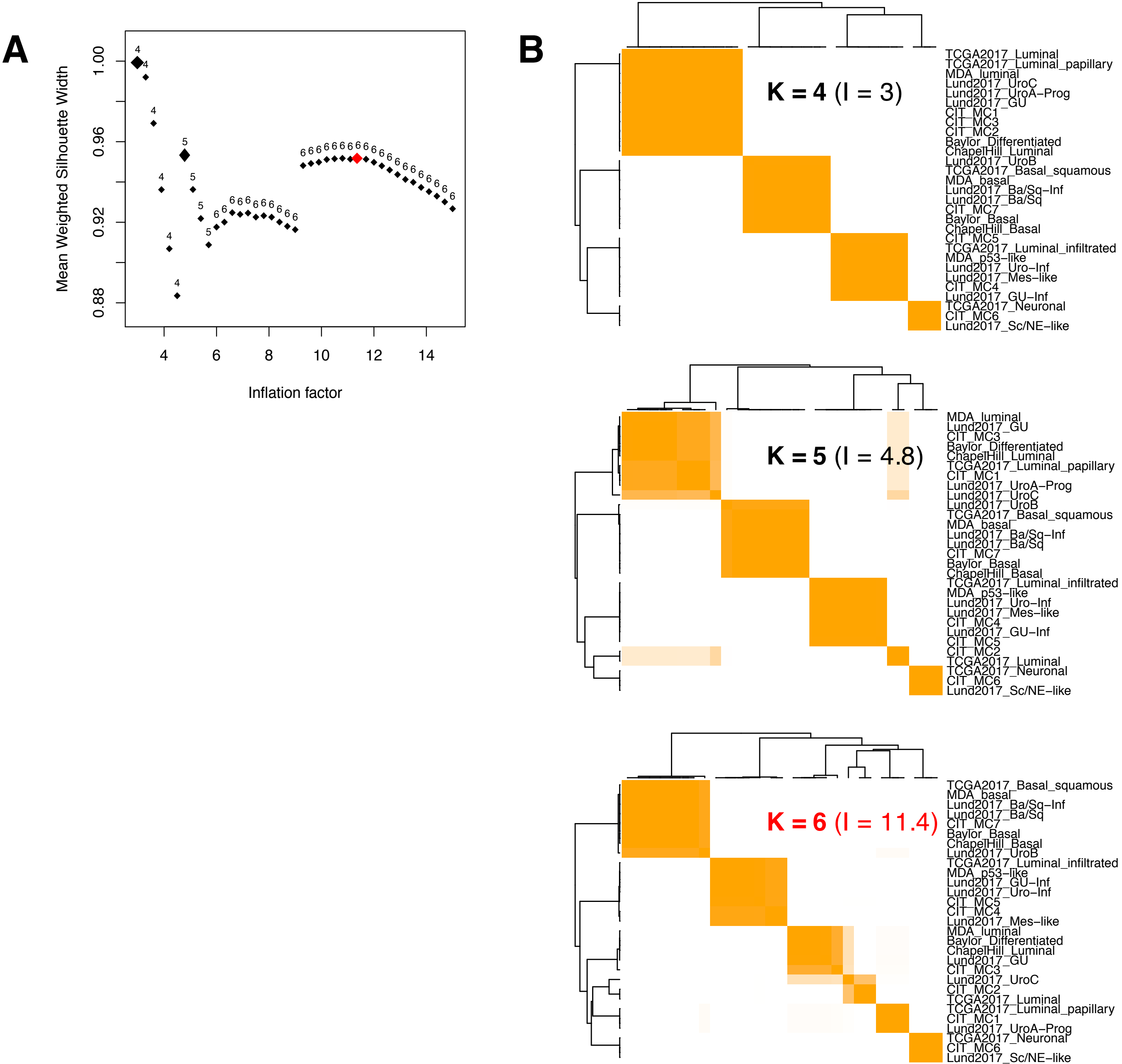

### Figure S3

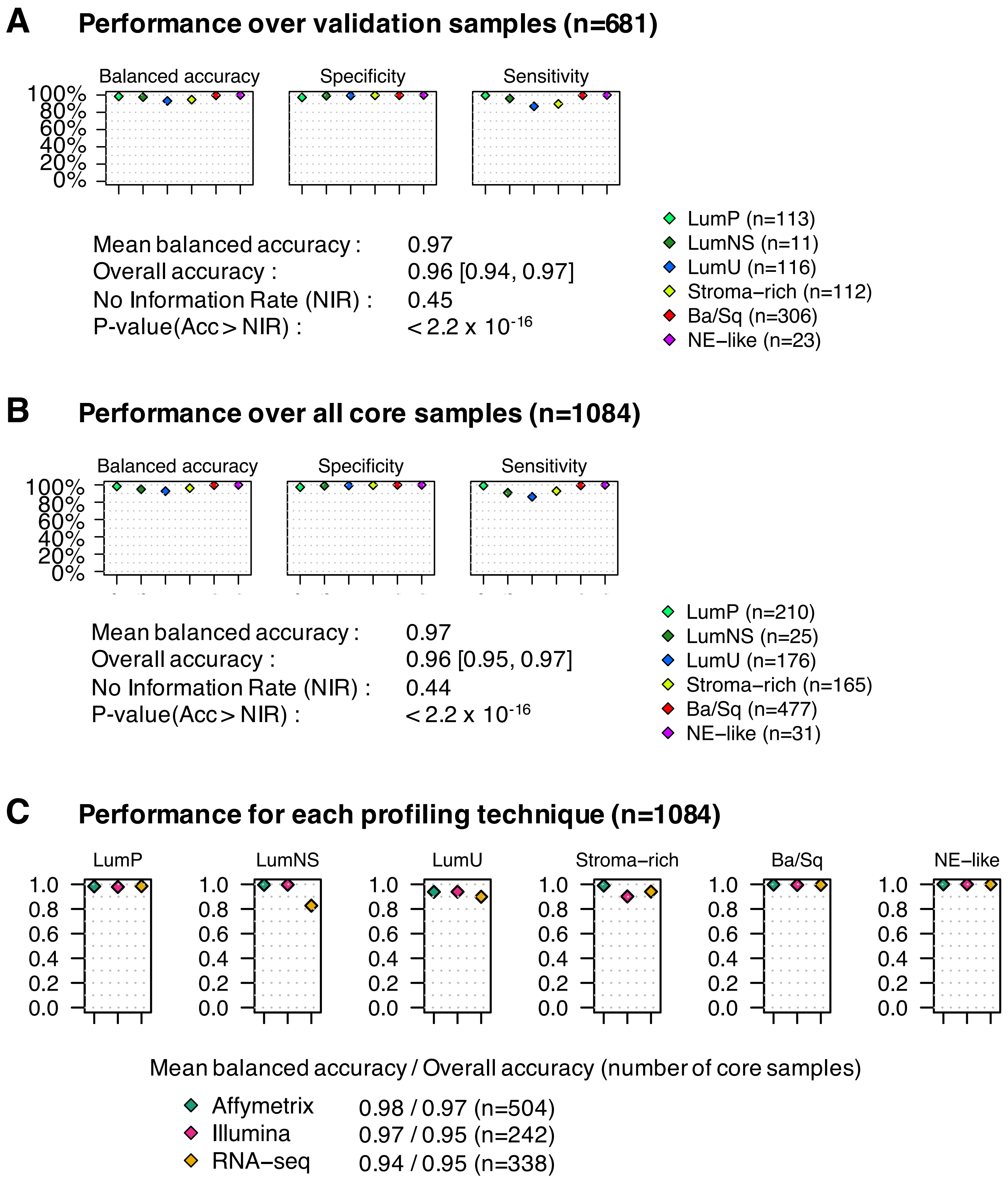

### Figure S4

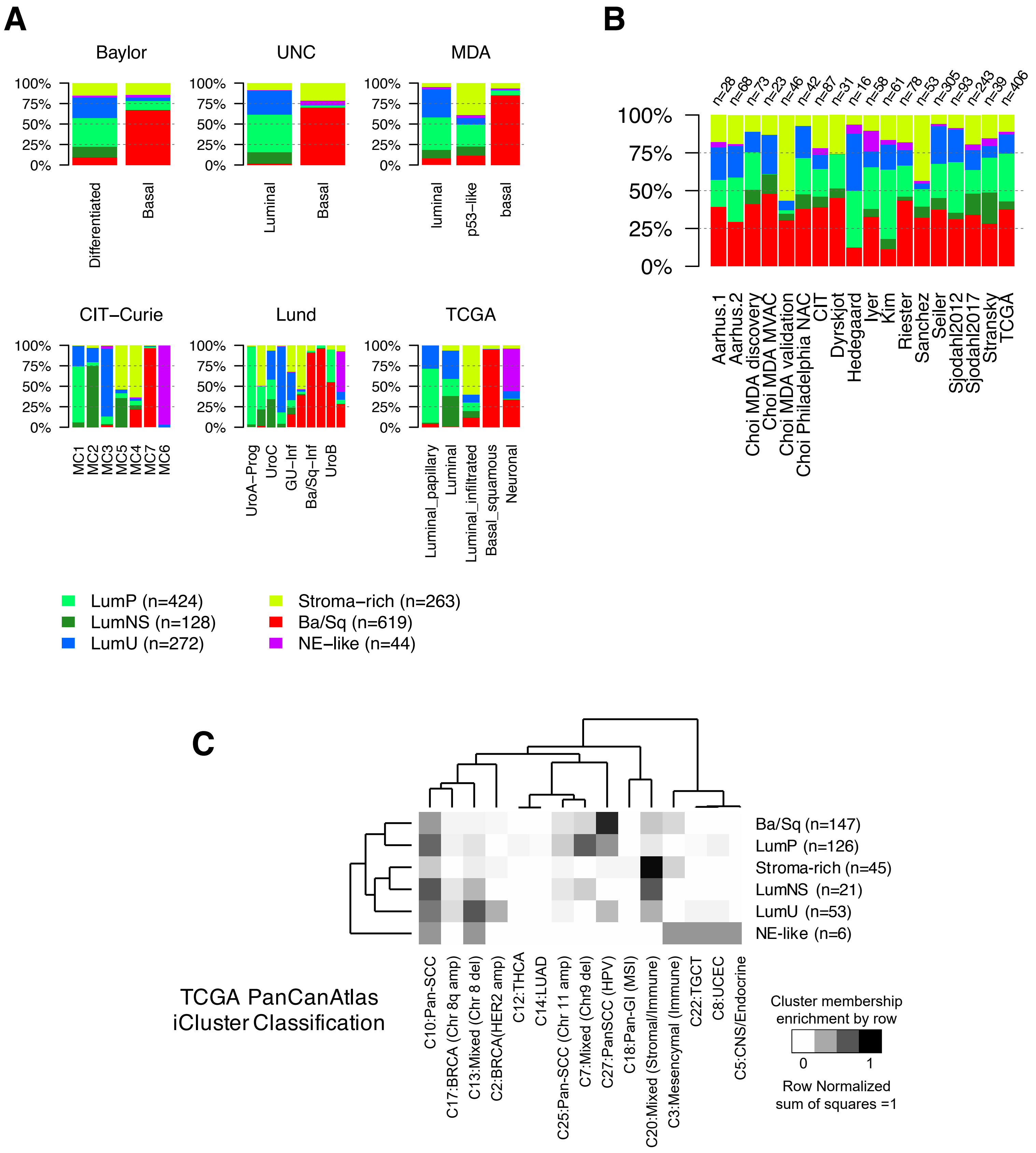

### Figure S5

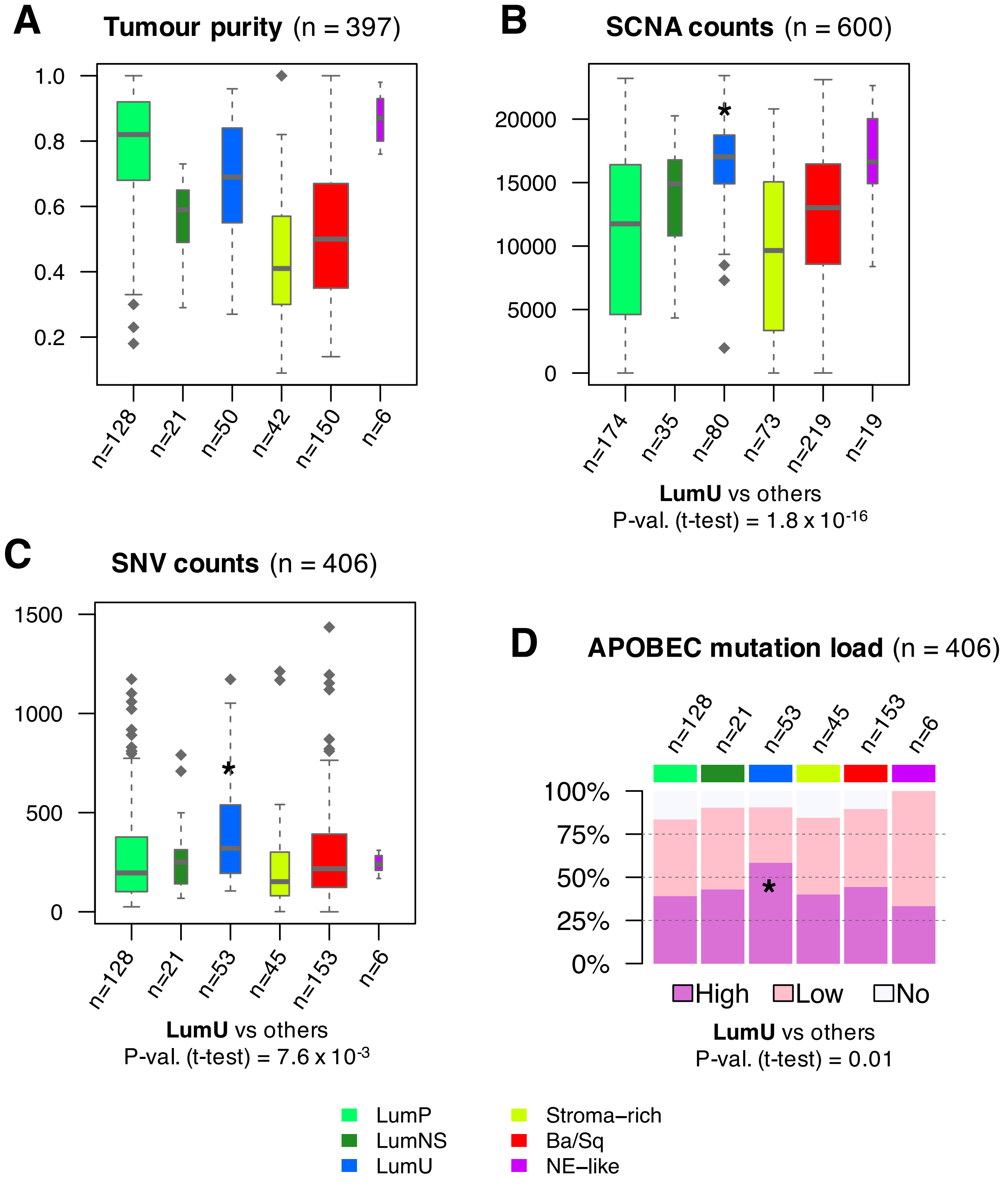

### Figure S6

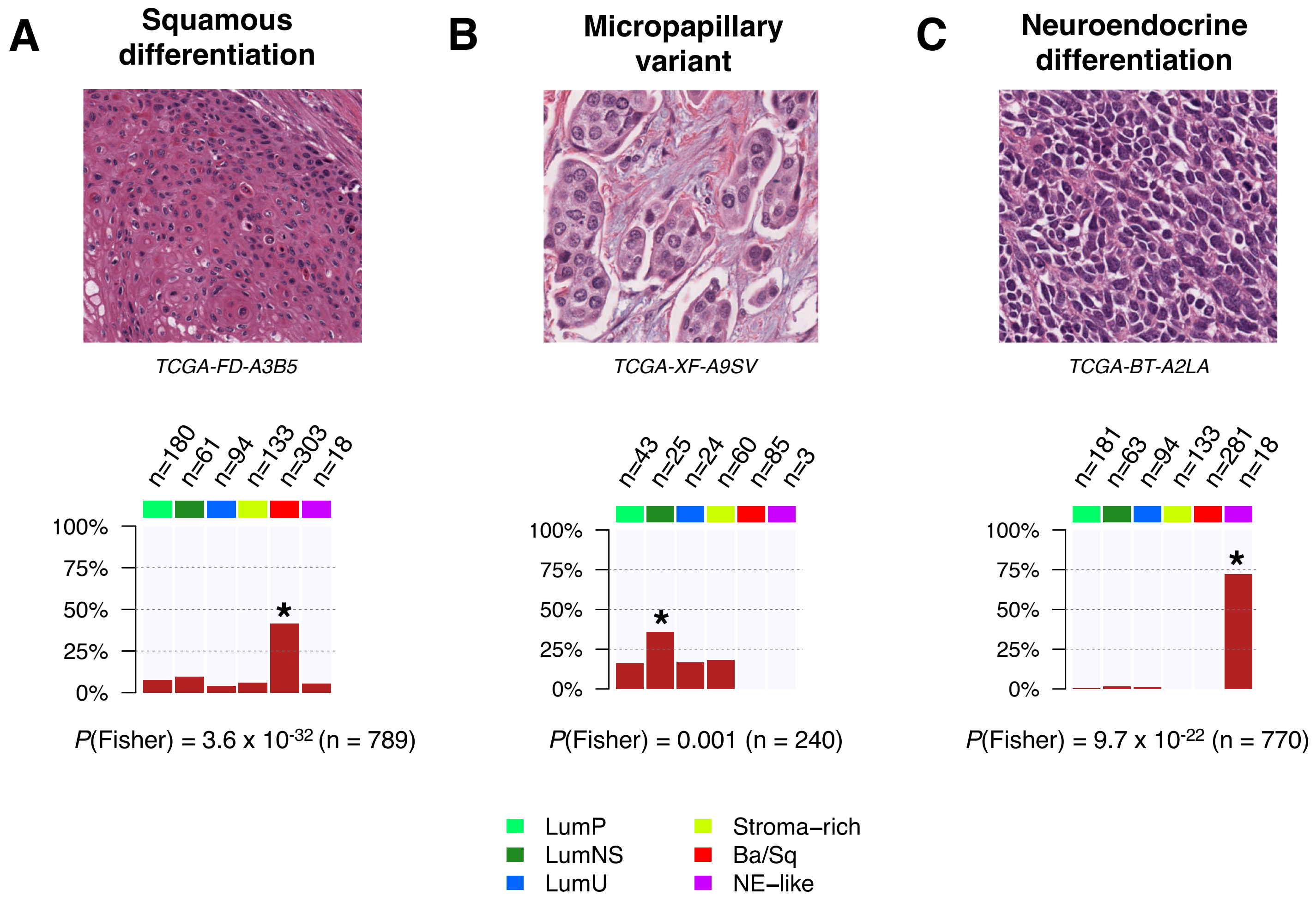

### Figure S7

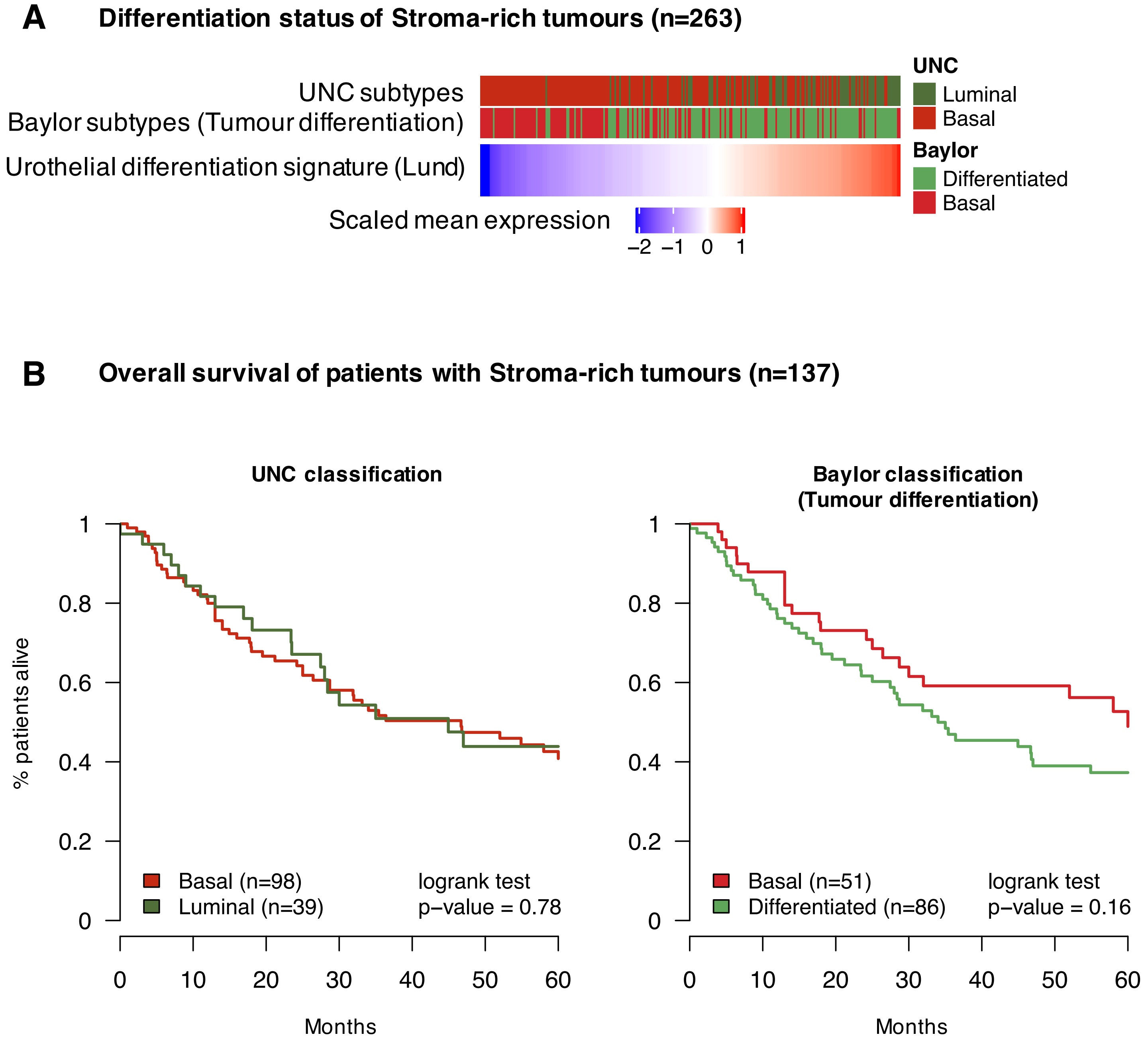
