## Supplementary Note for "A consensus molecular classification of muscle-invasive bladder cancer"

**Supplementary Note. Related to Figure 2.**

**Choice of names for the consensus molecular classes**

Differentiation states, as assessed by transcriptomic markers, were key determinants in choosing consensus class names, and the first part of each name indicates the differentiation signature that a class is associated with: Luminal, Basal, or Neuroendocrine. The one exception to this, the Stroma-rich class, is discussed below.

We discussed numbering each consensus class (1 to 6), or each luminal class (i.e. Lum 1 to 3), but chose names rather than numbers, and preferred simpler names over longer names that could be more precise.

We declined to name classes using tumor aggressiveness, as our understanding of aggressiveness may change with future studies involving larger cohorts, and with new treatments.

**Luminal Papillary (LumP)**

We named the LumP class from the papillary morphology that was frequent (~60%) in this class of tumors, though we note that this morphological pattern was also observed in LumNS (42%) and LumU (31%) tumors. Although LumP muscle-invasive (MI) tumors may have arisen as a progression from non-muscle invasive (NMI) papillary tumors, we caution that the LumP MIBC class name should not be confused with NMI papillary tumors.

Because more than 50% of these tumors activate FGFR3 through either mutation, fusion, or genomic amplification, we also considered “Luminal FGFR3”. However, FGFR3 may not be an oncogenic driver for all tumors of this class, and mutated-FGFR3 tumors are also found in other consensus classes.

**Luminal Non-Specified (LumNS)**

This class represents only 8% of the 1750 MIBC tumors analyzed. Although it is enriched with micropapillary histological variants, and is significantly associated with carcinoma in situ, we had too few LumNS cases to confidently base a consensus class name on either association. Future data and studies may support associating this consensus class with genomic or histological properties; until then, we use “Non-Specified”.

We note that this subtype corresponds well with the Lund UroC subtype, which has very distinct genomic properties^1^.

We considered, but did not retain, the use of the single-word “Luminal”, since we use this to refer to all tumors with luminal differentiation: LumP, LumNS, LumU, and a subset of stroma-rich tumors.

**Luminal Unstable (LumU)**

The name “unstable” refers to the “genomically unstable” class name in the Lund molecular taxonomy, and our LumU consensus class includes all tumors classified as “genomically unstable” by the Lund classifier. LumU tumors display several features of genomic instability, such as higher overall and APOBEC mutation burdens, and more somatic copy number alterations.

We preferred the shorter “Luminal Unstable” over the longer, more precise, “Luminal Genomically Unstable”. We considered “Luminal GU”, but did not retain it, because “GU” could be mistaken as an abbreviation for Genitourinary.

**Stroma-rich**

While we prefer class names that relate to tumor cell properties, this consensus class includes tumors that have diverse differentiation states and tumor cell phenotypes. It includes both luminal and non-luminal tumors. Given this, instead of using a name that indicates a differentiation state, we used a name that reflects the microenvironment: “Stroma-rich”. On a case-by-case basis, a tumor from this class may actually be phenotypically identical to “LumP”, “LumNS”, “LumU”, “Basal”, or “NE-like”, but will be in a “stroma-infiltrated” context, in which the infiltrating stromal cells are principally smooth muscle, then fibroblasts and myofibroblasts.

Because this class strongly overlapped with “Infiltrated” subtypes from TCGA and Lund classifications, we considered the name “Infiltrated”, but did not retain it, to prevent confusion between immune (lymphocytes) and stromal infiltration.

We also considered “Desmoplastic”, but judged “Stroma-rich” to be more accurate and easily understandable.

**Basal/Squamous (Ba/Sq)**

“Basal/Squamous” reflects the histological and molecular characteristics of the corresponding tumors, notably high levels of KRT14, KRT5/6, and a lack of GATA3 and FOXA1. The denomination “Basal/Squamous-like” was recommended by the current group of experts in 2015^2^. The “-like” extension was left out to reflect the true squamous differentiation that characterized a subgroup of these tumors. In addition, in pan-cancer analysis Hoadley et al.,2018), this class is highly correlated with cluster 27 (squamous cell carcinoma tumors (Figure S4C).

Although only 42% of histologically reviewed tumors within this class show squamous differentiation, this feature is very specific to these tumors, since 79% of tumors with squamous differentiation are classified within this group.

We discussed using the shorter names “Basal” or “Basal-like”, but suggest that these be used only informally.

**Neuroendocrine-like (NE-like)**

This consensus class is strongly associated with neuroendocrine histological variants: 75% of histologically reviewed tumors within this consensus class show neuroendocrine differentiation, and 81% of tumors with such histological features fall into this consensus class.

Although the TCGA “Neuronal” subtype is associated with this consensus class, we did not use this name because: 1) the NE-like consensus class is more strongly associated with neuroendocrine histology than the TCGA Neuronal subtype, and 2) bladder tumors are unrelated to neural cells.
